# Climate-smart prioritisation of tropical Key Biodiversity Areas for protection in response to widespread temperature novelty

**DOI:** 10.1101/2023.11.30.569430

**Authors:** Brittany T. Trew, Alexander C. Lees, David P. Edwards, Regan Early, Ilya M. D. Maclean

## Abstract

Key Biodiversity Areas (KBAs) are a cornerstone of 21^st^ century area-based conservation targets. In tropical KBAs, biodiversity is potentially at high risk from climate change, because most species reside within or beneath the canopy, where small increases in temperature can lead to novel climate regimes. We quantify novelty in temperature regimes by modelling hourly temperatures below the forest canopy across tropical KBAs between 1990 and 2019. We find that up to 66% of KBAs with tropical forest are likely to have transitioned to novel temperature regimes. Nevertheless, 34% of KBAs are providing refuge from novelty, 58% of which are not protected. By conducting the first pan-tropical analyses of changes in below-canopy temperatures, we identify KBAs that are acting as climate refugia and should be prioritised as candidates for expansion of the conservation network in response to the post-2020 Global Biodiversity Framework target to conserve 30% of land area by 2030.

## Introduction

Key biodiversity areas (KBAs) are sites of global importance for biodiversity in the face of an ongoing sixth mass extinction (Cowie, Bouchet, & Fontaine, 2022). They are identified following internationally recognised criteria that account for biodiversity metrics, such as the presence of globally threatened and/or range-restricted species (IUCN, 2016). KBAs foster species persistence (Butchart et al., 2015, 2012), and are a critical tool for an evidence-based approach to expand the global conservation network. The Post-2020 Global Biodiversity Framework includes a draft target to ensure that at least 30 per cent of land area globally is conserved by 2030 (CBD, 2020; Ward et al., 2020) and specifically identifies KBAs as a core priority for any expansion.

Tropical forests are global hotspots of terrestrial biodiversity (Barlow et al., 2018; Mittermeier et al., 2011). Yet, as well as increasing pressures from deforestation, fragmentation, and degradation (Vancutsem et al., 2021), there is an escalating threat from anthropogenic climate change. Below the canopy, tropical forests are climatically very stable, such that their biota has evolved under a narrow range of thermal conditions and may only be able to tolerate a small margin of warming above their thermal optima (Jirinec et al., 2022; Tewksbury et al., 2008; Trew & Maclean, 2021). As such, species here are at particularly high risk from novel climate conditions: climates with no historic analogues (Senior et al. 2019; Dobrowski et al., 2021; Trew et al., 2023). The ongoing transition of tropical forest environments to novel temperature regimes can easily precipitate changes in niche availability, favouring species with higher temperature affinity (Zellweger et al., 2020), and trigger changes to community composition through trophic cascades (Gilman et al., 2010; Lensing & Wise, 2006). As a result, shifting temperature regimes have the potential to undermine the effectiveness of tropical KBAs as a prioritisation tool for conservation strategy (Araújo et al., 2011).

Assessing the threat of climate change to tropical forest KBAs is a crucial step in prioritising effective protection or conservation initiatives on a site-by-site basis (Brown et al., 2022). However, thus far, we have had little to no understanding of how microclimates beneath the forest canopy - the conditions actually experienced by tropical forest organisms - are changing pan-tropically. We now have the ability to model microclimate conditions of the below-canopy tropical forest environment at high spatiotemporal resolutions, allowing us to understand where warming is already having the greatest impact on local below-canopy temperature regimes. Accordingly, we can use climate-smart prioritisation strategies to identify KBAs which are, thus far, not experiencing novel temperature regimes - currently acting as climate refugia - whose addition to the global conservation network using protected areas (PAs) or other effective conservation area-based conservation methods (OECMs) would greatly improve the future resilience of tropical biodiversity.

Here, we conduct the first global analyses of changes in below-canopy climatic conditions in tropical forests by integrating a recently developed mechanistic microclimate model (Maclean, 2023), with empirical temperature measurements and satellite-derived land-cover data, to quantify hourly below-canopy temperature at 5-km gridded resolution between 1990 and 2019. Across forests in the humid tropics, including tropical rain forest and tropical moist deciduous forest (hereafter tropical forests; *sensu* Vancutsem et al., (2021)), we derive an index of climate novelty from the fractional overlap in mean annual temperature between a baseline historical period (1990-2004) and the most recent historical period (2005-2019); results for six other temperature variables are presented in the supplementary information. Since tropical forests typically experience low temporal variability in temperature, minor changes can push climate conditions beyond a species’ normal thermal range. Consequently, we posit that evaluating temperature novelty is a more reliable measure of climate vulnerability than simply the magnitude of temperature changes (Foden et al., 2013). Ergo, the novelty index represents the fraction of years in the recent period in which mean annual temperatures lie outside the range of mean annual temperature experienced in recent history and identifies: (i) KBAs that are already highly threatened by shifting temperature regimes; and (ii) unprotected or partially protected KBAs that already provided refuge from shifting temperature regimes and should be priority areas for expansion of the global conservation network.

## Methods

Using a mechanistic microclimate model (Maclean 2023), we quantified hourly below-canopy climate conditions across the global tropics (-30 to 30°S; -109 to 180°E) between 1990 and 2019. The microclimate model was run in daily time increments and then hourly temperatures – at 0.05 m above the ground - were derived using the model’s interpolation methods, which infer hourly data from daily minima and maxima using the diurnal cycle in the ambient temperatures provided as inputs to the model.

The model is open-source and available as a documented R package on Github (Maclean, 2023). In summary, the following workflow is implemented. The model downscales hourly input climate-forcing data to the desired spatial resolution (in this case 5 km gridded resolution) using spatial interpolation and the application of an elevation- and humidity-dependent lapse rate correction. Temperature and water vapour at the desired height are then modelled mechanistically using principles of energy conservation, i.e., by assuming that components of the energy budget remain in balance, and by solving the energy budget for foliage temperature using the Penman-Monteith equation (Maclean and Klinges, 2021). Radiative energy is assumed to be influenced by slope, aspect, and canopy cover. Radiative fluxes through the canopy are estimated using a two-stream approximation model (Sellers, 1985). Sensible and latent heat fluxes are assumed to depend on wind speed, which in turn is attenuated vertically by canopy foliage using the method described in Harman and Finnigen (2007) and terrain-shelter adjusted using the method described in Ryan (1977). Latent heat fluxes are assumed additionally to depend on the stomatal conductance of leaves, which is quantified from the availability of photosynthetically active radiation using the method described in Kelliher et al. (1995). Ground heat fluxes are quantified from soil properties and from diurnal and annual cycles in temperature, using the method described in de Vries (1963) and also given in Campbell and Norman (2012). Air temperature is then derived from foliage temperatures using the localised near-field model described by Raupach (1989). Further details regarding the climate and environmental parameters, including canopy cover, driving the model are described in the supplementary methods. Validation of the modelled below-canopy temperatures across the global tropics was conducted and described in Trew et al., (2023).

The hourly modelled below-canopy climate conditions were used to calculate the annual bioclimatic variables detailed in Fick et al., (2017), namely: (1) mean annual temperature; (2) mean diurnal temperature range; (3) isothermality (diurnal range / annual range x 100); (4) seasonality; (5) maximum temperature of the warmest month; (6) minimum temperature of the coldest month; and (7) annual temperature range. The annual bioclimatic variables were split into a baseline historical time (1990 to 2004) and the most recent time (2005 to 2019). For each grid cell, we then derived an index of novelty for each variable from the fractional overlap in mean annual temperatures between a recent historical baseline time (1990 to 2004) and the most recent available time (2005 to 2019), calculated as one minus the proportional overlap between mean annual temperatures in the baseline and recent time periods. This novelty index represents the fraction of years in the recent period in which mean annual temperatures lie outside the range of mean annual temperatures that occurred in the historical baseline, whereby locations with a novelty index closer to 1 are those with no recent temperature analogue. Here, we have presented results for mean annual temperature, with results for the six other temperature variables in the supplementary information.

Global KBA boundaries (Birdlife International, 2022) were filtered to include KBAs that held at least one 5 km gridded cell of tropical forest (n = 2,663), including undisturbed and degraded tropical forest in 2019 as defined by Vancutsem et al., (2021), a detailed definition of which can be found in the supplementary methods. For each temperature variable, the mean fractional novelty occurring recently (2005 to 2019) was calculated by KBA and weighted by area. We define relatively novel temperature regimes as KBAs with >0.4 fractional mean novelty in the temperature variable when compared to the historic baseline and relatively stable temperature regimes as those with <0.4 fractional mean novelty in the temperature variable when compared to the historic baseline. Global PA boundaries were sourced from the World Database on Protected Areas (Protected Planet, 2021) and cleaned as per the standard protocol using the *wdpa* package (Hanson, 2022) in R (R Core, 2022). Unprotected tropical forest KBAs were identified by intersecting KBA and PA boundaries to calculate the percentage coverage of formal protection.

## Results

### Tropical KBAs are already highly threatened by shifting temperature regimes

Approximately 66% of KBAs holding tropical forests have recently transitioned to novel mean annual temperature regimes (>0.4 mean fractional novelty), with the remainder experiencing relatively stable temperature regimes over the last three decades. The proportion of KBAs in Africa and Latin America with relatively novel temperatures was particularly high (72% and 59%, respectively), while fewer KBAs across Asia and Oceania are shifting to relatively novel temperature regimes (49%).

There are KBAs across Latin America (2.9%) and a small number in Asia and Oceania (0.4%) that have recently transitioned to almost entirely novel temperature regimes (>0.8 mean fractional novelty). In Latin America, these KBAs were all located in Ecuador, Colombia, Venezuela, or Panama (Figure 1), with the Tropical Andes particularly affected by recent novel mean annual temperatures, including the Cayambe-Coca National Park (0.83) and Kutukú-Shaimi Protection Forest (0.84) of Ecuador. Across Asia, KBAs experiencing strong shifts in mean annual temperatures were predominantly located across Indonesia and the Philippines (Figure 3), including the Mt. Agtuuganon and Mt. Pasian KBA (0.86) and the indigenous territory of Pangasananan (0.85). We found no KBAs in Africa experiencing almost entirely novel mean annual temperature regimes, although some KBAs experienced strong shifts in recent temperature regimes, including the Gueoule and Glo Mountain Forest Reserves and Mt. Nimba Strict Nature Reserve (0.76 and 0.74, respectively), both located across Côte d’Ivoire and Guinea (Supplementary table S1). KBAs in the Central Congo basin moist forests in Democratic Republic of Congo have also transitioned to novel temperature regimes, including Africa’s largest tropical rainforest reserve: Salonga National Park (0.70).

**Figure 1.**
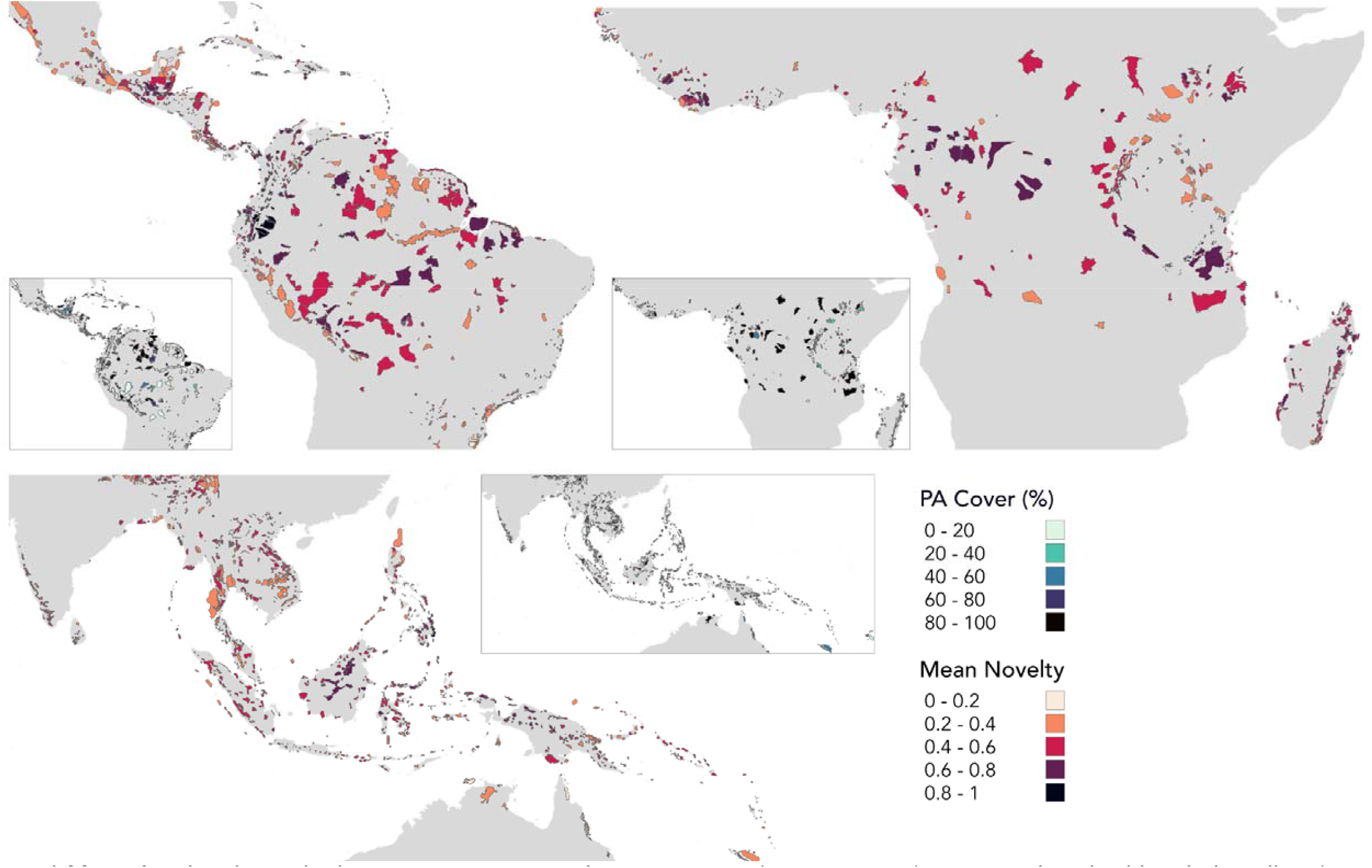
Mean fractional novelty in recent mean annual temperatures (2005 to 2019) compared to the historic baseline (1990 to 2004) across KBAs containing tropical forests in Latin America (n = 867), Africa (n = 395) and across Asia and Oceania (n = 1,262): 0-1, where 1 indicates entirely novel mean annual temperature regimes in 2005 - 2019. Individual inset maps show protected area coverage (%) for each KBA.

### Many tropical KBAs are acting as climate refugia but they lack protection

In Latin America, approximately 40% of KBAs experienced recent mean annual temperatures similar to the historic baseline, many of which were located in the Peruvian Yungas, Mexican Yucatan, and Guianan Shields. Latin America had the highest number of KBAs (0.06%) that experienced almost entirely un-novel mean annual temperature regimes and these were predominantly located in the coastal Atlantic Forest of Brazil. However, only 16% of climatically stable KBAs benefit from PA coverage over at least half their area and 6% do not benefit from any PA coverage (Figure 2). For example, the Sierra Madre Occidental Canyon Corridor in Northern Mexico (0.36) has no PA coverage (Figure 3). Of those KBAs with at least 80% PA coverage, 37% did not experience significant shifts in mean annual temperature regimes, including the UNESCO designated Iguazú National Park spanning Brazil and Argentina (0.27), part of the Atlantic Forest biome, and La Tigra National Park in Honduras (0.31).

**Figure 2.**
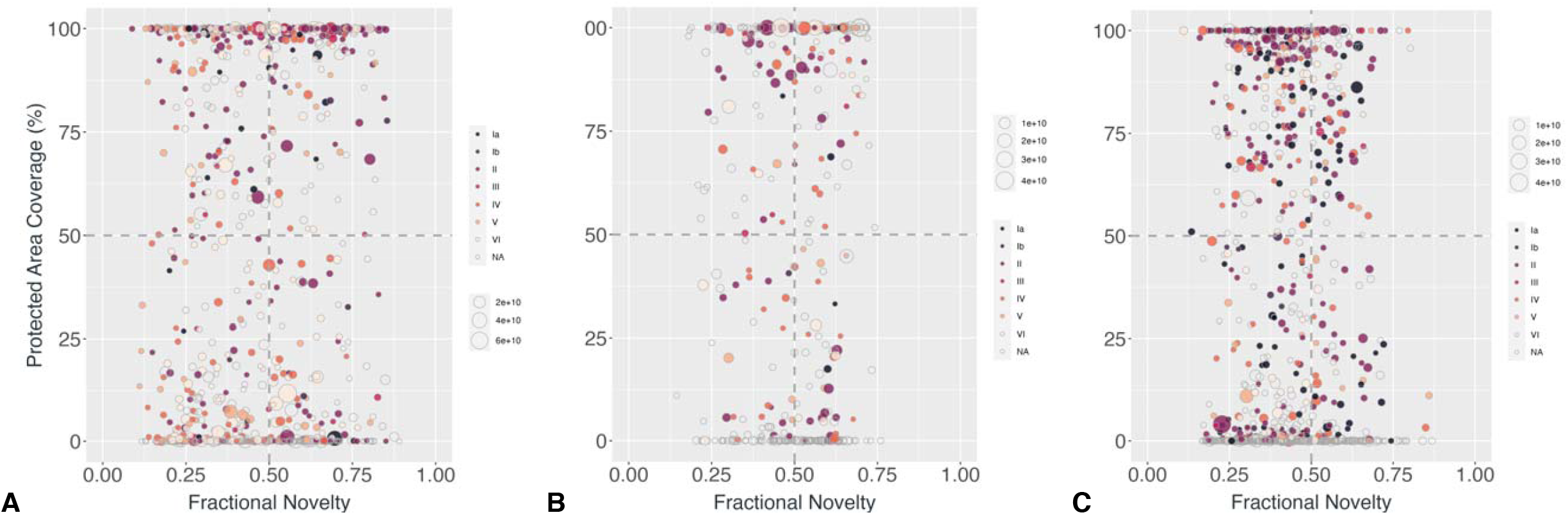
Each point represents a Key Biodiversity Area (KBA) containing tropical forest in (A) Latin America, (B) Africa, (C) Asia and Oceania and the relationship of the mean novelty in recent mean annual temperatures experienced by said Key Biodiversity Area and the formal protected area coverage (%) of the KBA. Point colours correspond to the IUCN category of the protected area and the size of each point corresponds to the geographical size of the KBA (m^2^). KBAs with low protected area coverage and low mean novelty in mean annual temperature should be prioritised for conservation planning.

**Figure 3.**
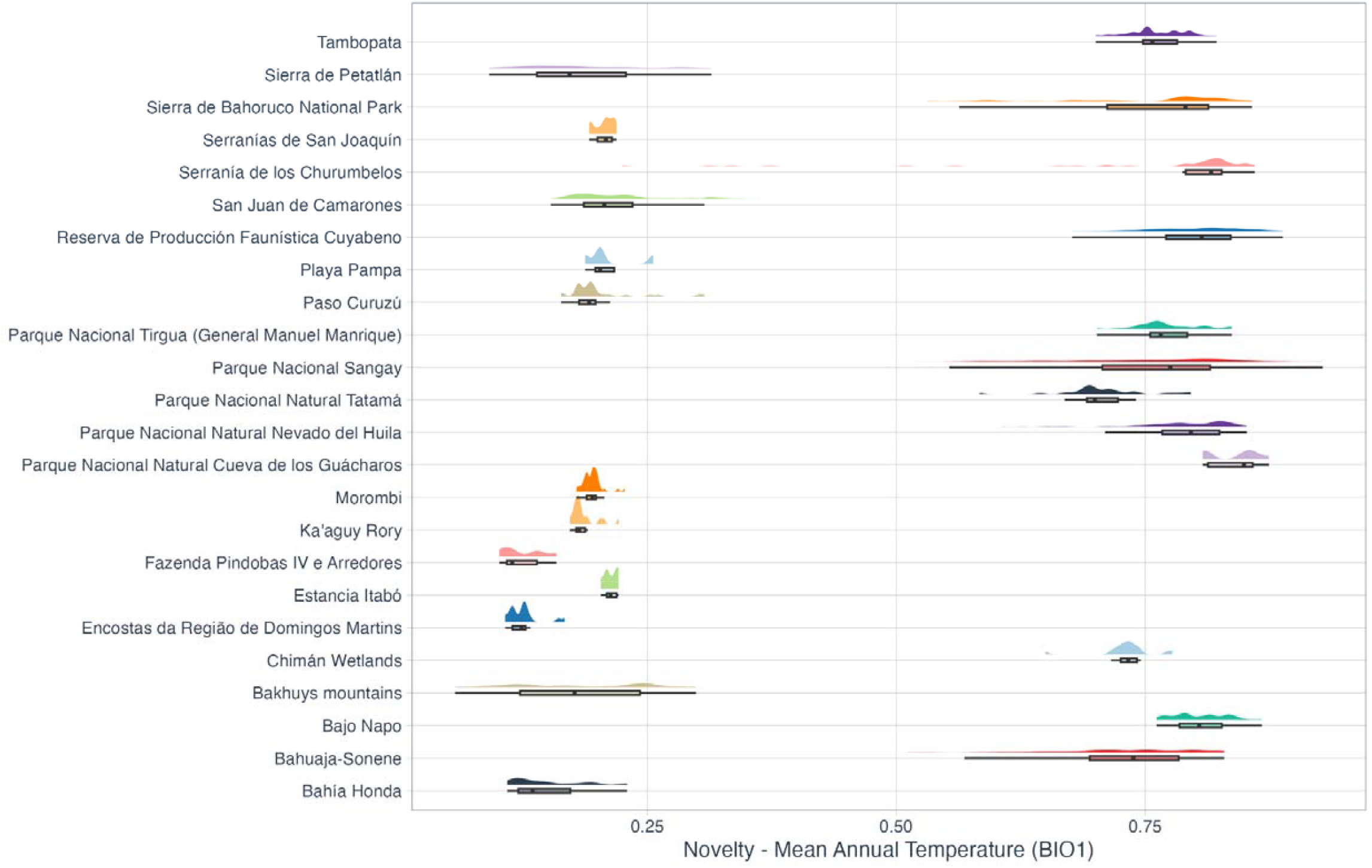
Fractional novelty in recent mean annual temperatures (2005 to 2019) compared to the historic baseline (1990 to 2004) for the 12 KBAs in **Latin America** containing tropical forest with at least 99% protected area coverage and the highest recent mean fractional novelty in mean annual temperature and the 12 KBAs in Latin America containing tropical forest with the lowest recent mean fractional novelty in mean annual temperature and no protected area coverage. Novelty is measured between 0-1, where 1 indicates entirely novel mean annual temperature regimes in 2005 - 2019.

Across Africa, far fewer KBAs experienced recent mean annual temperatures similar to the historic baseline (24%); these were mainly located in the Western Congo Basin, the East African montane forests and the highland forests across Equatorial Guinea and Cameroon. Only four KBAs experienced almost entirely stable mean annual temperature regimes, including the Rio Pongo and Iles Tristao KBAs and Ramsar sites in Guinea (both 0.18) and forest reserves in Uganda (Mount Kadam - 0.18) and Kenya (Kitale West - 0.14). Moreover, 13% of KBAs which have not shifted to novel mean annual temperature regimes were found to have PA coverage over at least half their area and 4% do not benefit from any PA coverage. For example, the South Nguruman KBA (0.20), forming the western wall of the Rift Valley in Kenya, has no PA coverage. (Figure 4). Of those KBAs with at least 80% PA coverage, 20% did not experience significant shifts in mean annual temperature regimes, including the Western Area Peninsula Forest National Park (0.24) in Sierra Leone, an important remnant of West African rainforest.

**Figure 4.**
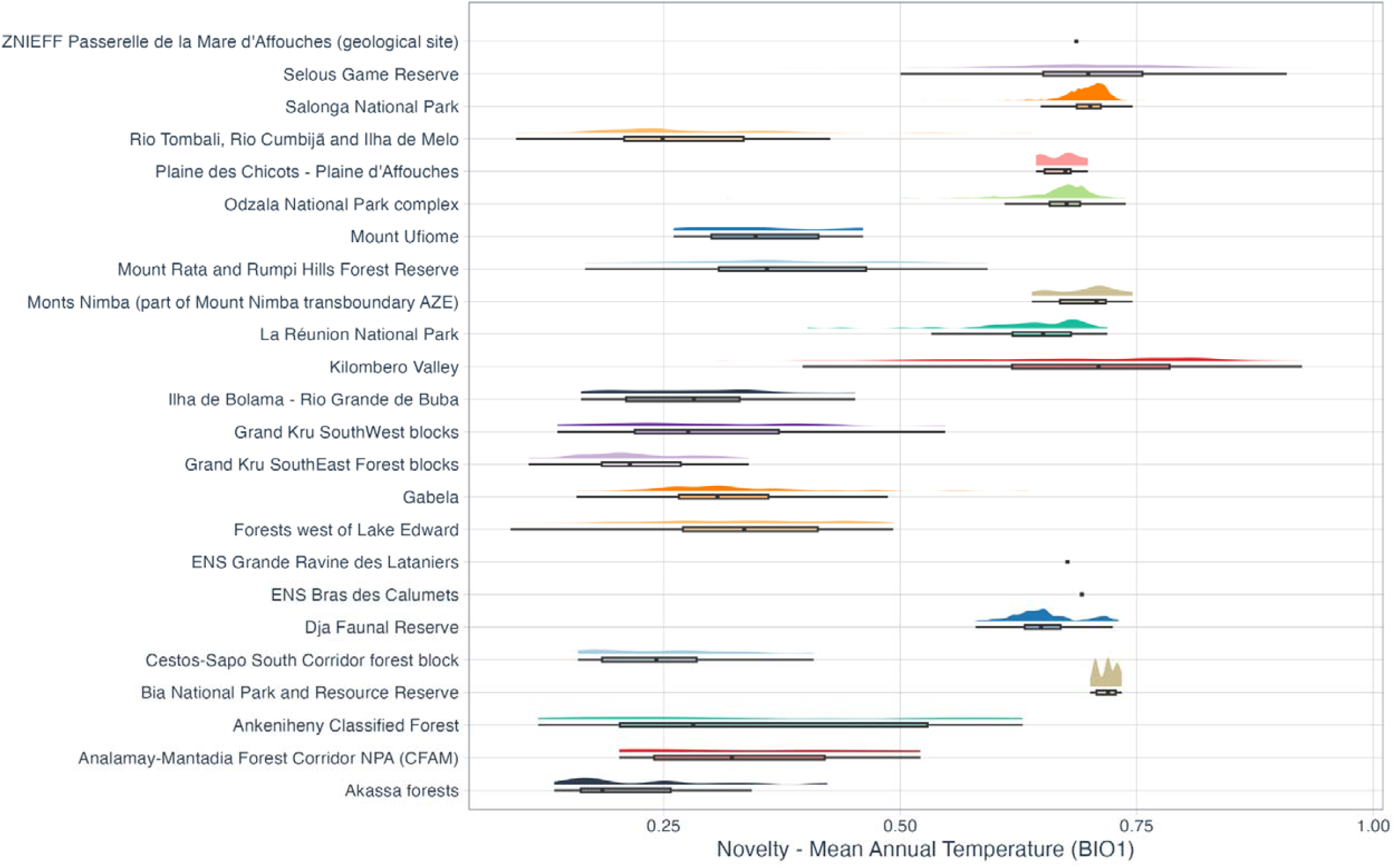
Fractional novelty in recent mean annual temperatures (2005 to 2019) compared to the historic baseline (1990 to 2004) for the 12 KBAs in **Africa** containing tropical forest with at least 99% protected area coverage and the highest recent mean fractional novelty in mean annual temperature and the 12 KBAs in Africa containing tropical forest with the lowest recent mean fractional novelty in mean annual temperature and no protected area coverage. Novelty is measured between 0-1, where 1 indicates entirely novel mean annual temperature regimes in 2005 - 2019.

Of the KBAs in Asia and Oceania, 46% experienced recent mean annual temperatures similar to the historic baseline. These KBAs were predominantly located across Southern Papua New Guinea, Central mainland Malaysia, and North-East Borneo. Only 0.02% of KBAs across Asia and Oceania experienced almost entirely stable mean annual temperature regimes (<0.2 mean fractional novelty). For instance, KBAs in Northern Australia’s tropical forests experienced some of the least novel temperature regimes globally, including Daintree Rainforest (0.11) and Wooroonooran National Park (0.17). However, a considerable number of KBAs experiencing low novelty in recent mean annual temperatures also lacked PA coverage (Figure 2). The Tiwi islands in North-West Australia experienced relatively un-novel temperatures over (0.19 fractional novelty) without benefiting from any formal PA coverage (Figure 5). Indeed, only 12% of KBAs which haven’t shifted to novel temperature regimes across Asia and Oceania have PA coverage over at least half of their extent and 23% did not benefit from any PA coverage. Of those KBAs with at least 80% PA coverage, 48% did not experience significant shifts in mean annual temperature regimes, including the world’s largest tiger reserve, the Hukaung Valley Wildlife Sanctuary KBA in the Northern Forest Complex of Myanmar (Figure 5).

**Figure 5.**
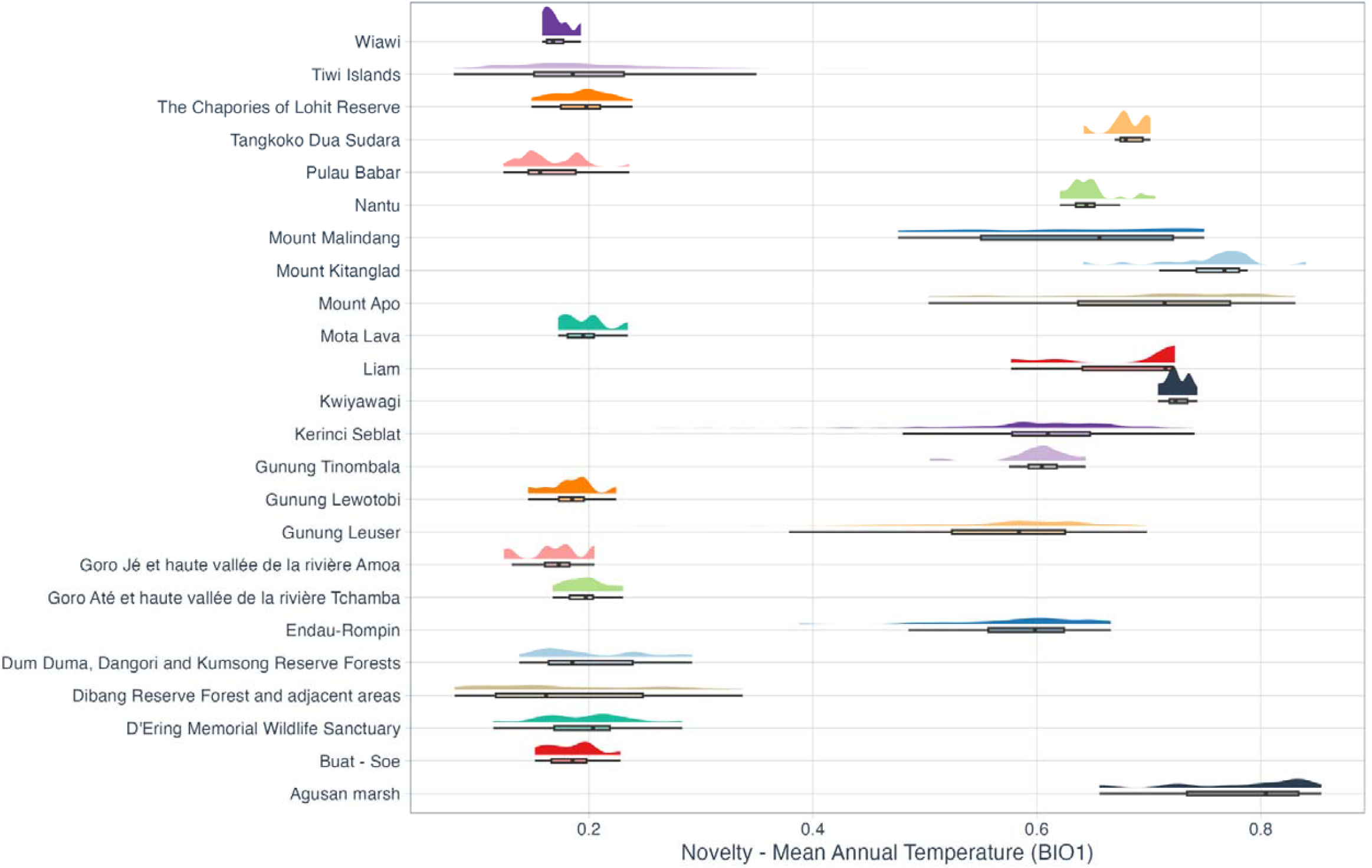
Fractional novelty in recent mean annual temperatures (2005 to 2019) compared to the historic baseline (1990 to 2004) for the 12 KBAs in **Asia and Oceania** containing tropical forest with at least 99% protected area coverage and the highest recent mean fractional novelty in mean annual temperature and the 12 KBAs in Asia and Oceania containing tropical forest with the lowest recent mean fractional novelty in mean annual temperature and no protected area coverage. Novelty is measured between 0-1, where 1 indicates entirely novel mean annual temperature regimes in 2005 - 2019.

## Discussion

Global conservation efforts would be increasingly effective if prioritisation methods considered the potential impact of ongoing climate changes. Ongoing shifts from climate regime baselines are not uniform across the globe and the impact of these shifts depends, in part, on the prehistoric variability of climate (Trew & Maclean, 2021) which varies geographically and vertically in space (De Frenne et al., 2021, Lembrechts et al., 2018). As a result of naturally lower climate variability below the forest canopy, tropical forests are now subject to increasing threat from novel climates (IPCC, 2021; Trew et al., 2023). This is particularly concerning as tropical forests host the majority of biodiversity worldwide (Gardner et al., 2010; Pillay et al., 2022) and KBAs here hold particular conservation value, having been identified on the basis of the biodiversity they host as well as holding some of the largest areas of high integrity forest (Crowe et al., 2023). Thus, it is crucial that KBAs which are so far providing refuge from shifting temperature regimes and those already at risk are identified to prioritise climate-smart decision-making. To achieve this, we quantified the recent novelty of below-canopy temperatures across KBAs with tropical forest between 1990 and 2019 relative to a historic baseline. As well as highlighting a large proportion of KBAs already impacted by novel temperature regimes, many of which are internationally important national parks and indigenous lands (Garnett et al., 2018), we have identified substantial numbers of KBAs that could benefit from expansion of the global conservation network using protected areas, OECMs or other approaches.

### Biodiversity in KBAs experiencing climate novelty is at high risk

Conservation schemes in regions affected by shifts to novel temperature regimes will need to explicitly consider climate-driven changes in biodiversity patterns (Dowbrowski et al., 2021). Exemplars of KBAs already experiencing novel temperature regimes are those within the Yasuní Biosphere Reserve—a designated Ecuadorian UNESCO World Heritage Site with some of the highest biodiversity per square metre globally (Bass et al., 2010)—which have experienced some of the strongest shifts in recent temperature regimes pan-tropically. KBAs within both lowland PAs like Salonga National Park in the Democratic Republic of Congo and PAs covering mountainous forests such as the Mount Nimba Strict Nature Reserve have also recently experienced strong shifts to novel temperature regimes. Species inhabiting lowland areas will struggle to track their environmental niche due to an absence of elevational gradients (Trew and Maclean, 2021). Equally, many species inhabiting mountain regions such as Mount Nimba —a UNESCO World Heritage Site in West Africa that encompasses 1,752 metres in elevation— could be highly vulnerable to temperature changes. Species here are likely to have narrow ranges and limited dispersal ability. Consequently, novel temperature regimes may mean they lose access to their climate envelope and are outcompeted by downslope taxa (Enquist et al., 2002; Laurance et al., 2011).

To mitigate anticipated biodiversity loss within KBAs experiencing high rates of local climate change, it is paramount that distant wealth-related drivers of deforestation, degradation and climate change are addressed (Carmenta et al., 2023). Locally, large-scale forest restoration programmes are needed (Gillson et al., 2013) within and outside of KBAs to connect forest fragments and promote climate connectivity, as well as the overall size and interior (i.e. non-edge effected) of forest (Gonzalez del Pliego et al., 2016; Strassburg et al., 2020). Such restoration efforts and any expansion of a conservation network must be undertaken in a socially just manner, preferably with local organisations representing people living in and managing tropical forest landscapes (Fleischman et al. 2022).

### Prioritising climate-smart KBAs for expansion of the global conservation network

Fortunately, there are considerable numbers of unprotected and partially protected KBAs, highlighted here, that provide refuge from novel temperature regimes and are prime candidates for expanding conservation programs. For example, the Central Suriname Nature Reserve - a UNESCO World Heritage Site - covers multiple KBAs and has a large elevational range (over 1,200 metres); safeguarding multiple habitats that support exceptional levels of biodiversity and act as a buffer against climate change impacts. However, as is the case for many tropical PAs (Mascia & Pailler, 2011; Leberger et al., 2020), there is very little on-the-ground management capacity to address intensifying threats from human activities such as nearby mining and logging (IUCN’s Conservation Outlook Assessment, 2020). Pressures on biodiversity can accumulate if protections for KBAs are inadequate or under-resourced, as climate change can also interact with and act as a multiplier of anthropogenic threats like habitat fragmentation and degradation (Bowler et al. 2019).

For ambitious area-based conservation targets like the Post-2020 Global Biodiversity Framework’s *30by30* to be effective, expansion of the global conservation network needs to urgently consider the spatiotemporal patterns of climate change and prioritise the protection of climate-smart KBAs. Pantropically, this means those KBAs which have experienced low novelty in recent temperature regimes are among the highest priorities for assessing required conservation actions to target drivers of forest loss and degradation, especially in locations where pressure is highest, via a combination of legal protection (Roberts et al., 2020), carbon payments (Crossman et al., 2011), empowering indigenous communities (Sze et al., 2022), or other effective area-based conservation methods (Dudley et al., 2018). Moreover, to ensure an adequately representative sample of pantropical biodiversity is protected, conservation programs should consider how climate-smart KBAs can be prioritised across biogeographical realms. In this way, global conservation efforts would be increasingly effective if prioritisation considered ongoing climate change impacts to improve the future resilience of global tropical biodiversity.

## Acknowledgements

IM was supported by NERC (grants NE/W006618/1 and NE/X015262/1).

## Data Availability Statement

Global hourly climate data is available from https://cds.climate.copernicus.eu/. Environmental parameters include: (a) leaf area index & surface reflectance available from https://www.ncei.noaa.gov/data/avhrr-land-leaf-area-index-and-fapar/, (b) global habitat types available from https://www.esa-landcover-cci.org/, (c) vegetation height available from https://webmap.ornl.gov/ogc/, (d) soil types available from https://www.soilgrids.org, (e) digital elevation model available from: https://www.usgs.gov/centers/eros/science/usgs-eros-archive-digital-elevation-shuttle-radar-topography-mission-srtm-1. The microclimate model is freely available for download and adaptation via a GitHub repository: https://github.com/ilyamaclean/microclimf. The global tropical forest monitoring dataset is available from https://forobs.jrc.ec.europa.eu/TMF. Temperature records used for validation are available from the global SoilTemp dataset on request: https://www.soiltempproject.com/the-soiltemp-database/. Protected area shapefiles are freely available from the Protected Planet database: https://www.protectedplanet.net. Key Biodiversity Area shapefiles are available on request: www.keybiodiversityareas.org.

## Code Availability Statement

Code used for the novelty analysis is published online (10.5281/zenodo.8246818) with examples of the open access datasets needed to reproduce the results shown here (those listed in the data availability statement). The mechanistic microclimate model is freely available to use in the *microclimf* package for R: https://github.com/ilyamaclean/microclimf.

