## Supplementary Information for "Climate-smart prioritisation of tropical Key Biodiversity Areas for protection in response to widespread temperature novelty"

**1.0 Detailed Methods**

**1.1 Climate data and modelling**

Using a mechanistic microclimate model (Maclean and Klinges, 2023) we quantified hourly below-canopy climate conditions across the global tropics (-30 to 30°S; -109 to 180°E) between 1990 and 2019. The microclimate model was run in daily time increments and then hourly temperatures – at 0.05 m above the ground - were derived using the model's interpolation methods, which infer hourly data from daily minima and maxima using the diurnal cycle in the ambient temperatures provided as inputs to the model. Full details of the model are provided in Maclean and Klinges (2023), but in summary the following workflow is implemented. First, the model downscales hourly input climate-forcing data to the desired spatial resolution (in this case 5 km gridded resolution) using spatial interpolation and the application of an elevation and humidity-dependent lapse rate correction. Temperature and water vapour at the desired height are modelled mechanistically using principles of energy conservation, i.e. by assuming that components of the energy budget remain in balance, and by solving the energy budget to derive differences between near-ground and ambient temperature using the Penman-Monteith equation as described in Maclean and Klinges (2021). Radiative energy is assumed to be influenced by slope, aspect, and a leaf area index. Radiative fluxes through the canopy are estimated using the Sellers two-stream approximation model (Sellers, 1985). Sensible and latent heat fluxes are assumed to depend on wind speed, which in turn is attenuated vertically by canopy foliage using the method described in Raupach (1994). Wind speed is terrain-adjusted using the method described in Ryan (1977). Latent heat fluxes are assumed additionally to depend on the stomatal conductance of leaves, which is quantified from the availability of photosynthetically active radiation using the method described in Kelliher et al. (1995). Ground heat fluxes are quantified from canopy-soil temperature gradients, the latter contingent primarily on radiation absorbed by the ground using the method described in Campbell and Norman (2012).

The hourly climate-forcing data required to drive the microclimate model were obtained from the ERA5 fifth generation ECMWF atmospheric reanalysis of the global climate, using the single levels surface dataset (Hersbach et al., 2020) at a 0.25 degree gridded spatial resolution for the 30-year time. The ERA5 climate data assimilates past climate observations with climate model predictions to generate a series of climate variables for atmospheric, land-surface, and sea parameters. The following climate variables were extracted for the extent of the study area: (1) air temperature at 2 metres, (2) specific humidity at surface, (3) pressure at surface, (4) precipitation rate, (5) U wind at 10 metres, (6) V wind at 10 metres, (7) total cloud cover, (8) sky emissivity and (9) downward solar radiation, which was partitioned into direct and diffuse components using the method by Skartveit *et al. (*1998).

Additionally, the following environmental predictors were obtained to drive the microclimate model (i) annual habitat type, sourced from the European Space Agency Climate Change Initiative (ESA, 2017) at a gridded spatial resolution of 5 km; (ii) annual vegetation height, sourced from ORNL DAAC (Dubayah et al., 2020) at a gridded spatial resolution of 5 km; (iii) monthly plant area index, calculated as the sum of monthly leaf area index (LAI) and 20% of the monthly maximum LAI. Monthly LAI values were sourced from the National Oceanic and Atmospheric Administration (Vermote et al., 2014) and spatio-temporally aggregated to a gridded spatial resolution of 5 km, with missing values estimated from the LAI at the same location in other months using a locally-informed ’month effect accounting for seasonal cycles; (iv) monthly canopy and ground reflectance at a spatial resolution of 5 km gridded resolution, calculated by first deriving fractional canopy cover from surface albedo (Vermote et al., 2014) and monthly LAI values and then using the fractional canopy cover to partition surface albedo between ground and canopy; both steps used the *microclima* (Maclean, 2019) package for R 4.2 (R Core Team, 2022); (v) soil type, sourced at a gridded spatial resolution of 250 m from soilgrids.org (Hengl et al., 2017), which was then resampled to a gridded spatial resolution of 5 km using the nearest neighbour method; (vi) a digital elevation model, sourced from the U.S. Geological Survey (Danielson & Gesch, 2011) at a gridded spatial resolution of 7.5-arc-second and resampled to 5 km using a bilinear method; and (vii) a topographic wetness index at a gridded spatial resolution of 5 km, calculated by using the digital elevation model to derive flow accumulation. The microclimate model was validated against observations from temperature loggers across the global tropics in a previous study (see: Trew et al., 2023).

**1.2 Climate novelty analysis**

The hourly modelled below-canopy climate conditions were used to calculate the annual bioclimatic variables detailed in Fick et al., (2017), namely: (1) mean annual temperature, (2) mean diurnal temperature range, (3) isothermality (diurnal range / annual range x 100), (4) seasonality, (5) maximum temperature of the warmest month, (6) minimum temperature of the coldest month, and (7) annual temperature range. The annual bioclimatic variables were then split into a baseline historical time period (1990 to 2004) and the most recent time period (2005 to 2019). For each grid cell, we then derived an index of novelty for each variable from the fractional overlap in mean annual temperatures between a recent historical baseline time (1990 to 2004) and the most recent available time (2005 to 2019), calculated as: 1 minus the proportion of overlap between mean annual temperatures in the baseline time period and the recent period. This novelty index represents the fraction of years in the recent period in which mean annual temperatures lie outside the range of mean annual temperatures that occurred in the historical baseline. Locations with a novelty index closer to 1 are those with no recent temperature analogue, relative to the recent historical baseline. Here, we have presented results for mean annual temperature, with results for six other temperature variables available in the supplementary information.

To exclude forest where climate change could be amplified by interacting human activities like deforestation, novelty index values for each of the seven bioclimatic variables were extracted for the locations of tropical moist forest which were still undisturbed in 2019, as defined by Vancutsem et al., (2021) as all closed forests in the humid tropics including the tropical rainforest and the tropical moist deciduous forest without any disturbance (degradation or deforestation) and observed over the full observation period defined by the available Landsat data (1982 - 2019). The definition is not based on percentage of canopy cover and does not discriminate between primary and secondary growth tropical forest because there is no Landsat data available prior to 1982. However, it is likely that undisturbed tropical forest cover as estimated here is close to the true extent of primary tropical forest due to the amount of time that they have been undisturbed

**References.**

Campbell, G. S. & Norman, J. M. An introduction to environmental biophysics. Second edn, (Springer Science+Business Media, 1998).

Danielson, J.J., and Gesch, D.B., 2011, Global multi-resolution terrain elevation data 2010 (GMTED2010): U.S. Geological Survey Open-File Report 2011–1073, 26 p.

Dubayah, R.O., S.B. Luthcke, T.J. Sabaka, J.B. Nicholas, S. Preaux, and M.A. Hofton. 2021. GEDI L3 Gridded Land Surface Metrics, Version 2. ORNL DAAC, Oak Ridge, Tennessee, USA. doi:10.3334/ORNLDAAC/1952

ESA. Land Cover CCI Product User Guide Version 2. Tech. Rep. (2017). Available at: maps.elie.ucl.ac.be/CCI/viewer/download/ESACCI-LC-Ph2-PUGv2_2.0.pdf

Fick, S.E., Hijmans, R.J. (2017) WorldClim 2: new 1-km spatial resolution climate surfaces for global land areas. International Journal of Climatology 37, 4302-4315.

Hengl, T. et al. SoilGrids250m: Global gridded soil information based on machine learning. PLOS ONE 12, e0169748 (2017). [https://doi.org:10.1371/journal.pone.0169748](about:blank)

Hersbach, H. et al. The ERA5 global reanalysis. Quarterly Journal of the Royal Meteorological Society 146, 1999-2049 (2020). <https://doi.org/10.1002/qj.3803>

Kelliher, F. M., Leuning, R., Raupach, M. R. & Schulze, E. D. Maximum conductances for evaporation from global vegetation types. Agricultural and Forest Meteorology 73, 1-16 (1995). <https://doi.org/10.1016/0168-1923(94)02178-M>

Maclean, I. M. D. & Klinges, D. H. Microclimc: A mechanistic model of above, below and within-canopy microclimate. Ecological Modelling 451, 109567 (2021). https://doi.org/10.1016/j.ecolmodel.2021.109567

Maclean, I. M. D., Mosedale, J. R. & Bennie, J. J. Microclima: An r package for modelling meso- and microclimate. Methods in Ecology and Evolution 10, 280-290 (2019). [https://doi.org:10.1111/2041-210x.13093](about:blank)

Maclean, I. M. D., Klinges, D. H. (2023). Microclimf. Github repository. https://github.com/ilyamaclean/microclimf

R Core, T. (2022). R: A language and environment for statistical computing. . Vienna: R Foundation for Statistical Computing.

Raupach, M. R. Simplified expressions for vegetation roughness length and zero-plane displacement as functions of canopy height and area index. Boundary-Layer Meteorology 71, 211-216 (1994). https://doi.org:10.1007/BF00709229

Ryan, B. C. A Mathematical Model for Diagnosis and Prediction of Surface Winds in Mountainous Terrain. Journal of Applied Meteorology and Climatology 16, 571-584 (1977). [https://doi.org:10.1175/1520-0450](about:blank)

Sellers, P.J. (1985) Canopy reflectance, photosynthesis and transpiration. International Journal of Remote Sensing 6, 1335-1372.

Skartveit, A., Olseth, J.A., Tuft, M.E. (1998) An hourly diffuse fraction model with correction for variability and surface albedo. Solar Energy 63, 173-183.

Trew, B., Edwards, D., Lees, A., Klinges, D.H., Early, R., Svátek, M., Plichta, R., Matula, R., Okello, J., Niessner, A., Barthel, M., Six & Maclean, I. (2023) Novel climates are already widespread beneath the world’s tropical forest canopies. Research Square, Preprint.

Vancutsem, C., Achard, F., Pekel, J.F., Vieilledent, G., Carboni, S., Simonetti, D., Gallego, J., Aragão, L.E.O.C., Nasi, R. (2021) Long-term (1990–2019) monitoring of forest cover changes in the humid tropics. Science Advances 7, eabe1603.

Vermote, Eric; Justice, Chris; Csiszar, Ivan; Eidenshink, Jeff; Myneni, Ranga B.; Baret, Frederic; Masuoka, Ed; Wolfe, Robert E.; Claverie, Martin; NOAA CDR Program. (2014): NOAA Climate Data Record (CDR) of AVHRR Surface Reflectance, Version 4. NOAA National Centers for Environmental Information. doi:10.7289/V5TM782M.

**2.0 Supplementary Figures**

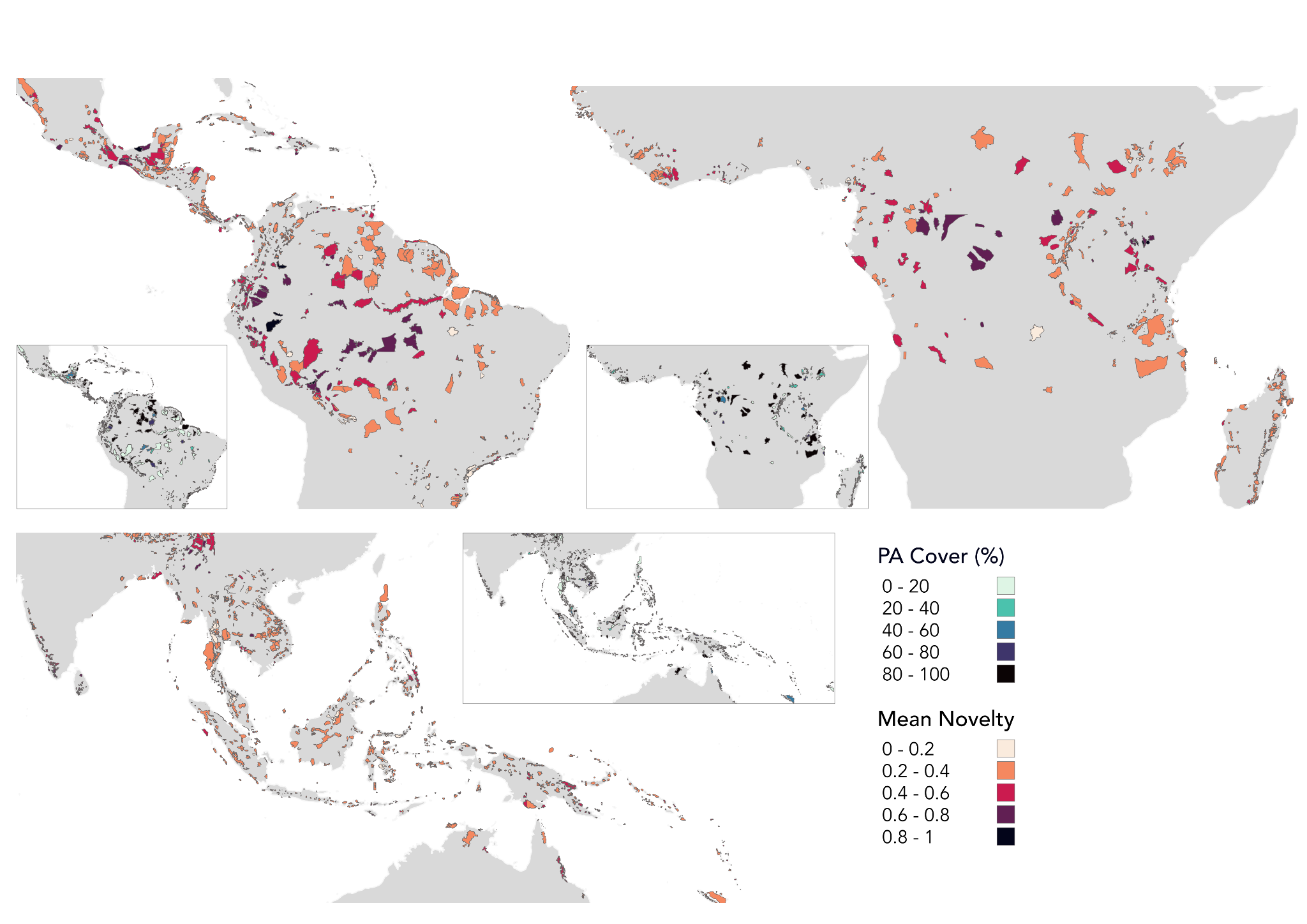

**Figure S2.1 |** Mean fractional novelty in recent **mean diurnal temperature range** (2005 to 2019) compared to the historic baseline (1990 to 2004) across KBAs containing tropical forests in Latin America (n = 867), Africa (n = 395) and across Asia and Oceania (n = 1,262): 0-1, where 1 indicates entirely novel mean annual temperature regimes in 2005 - 2019. Individual inset maps show protected area coverage (%) for each KBA.

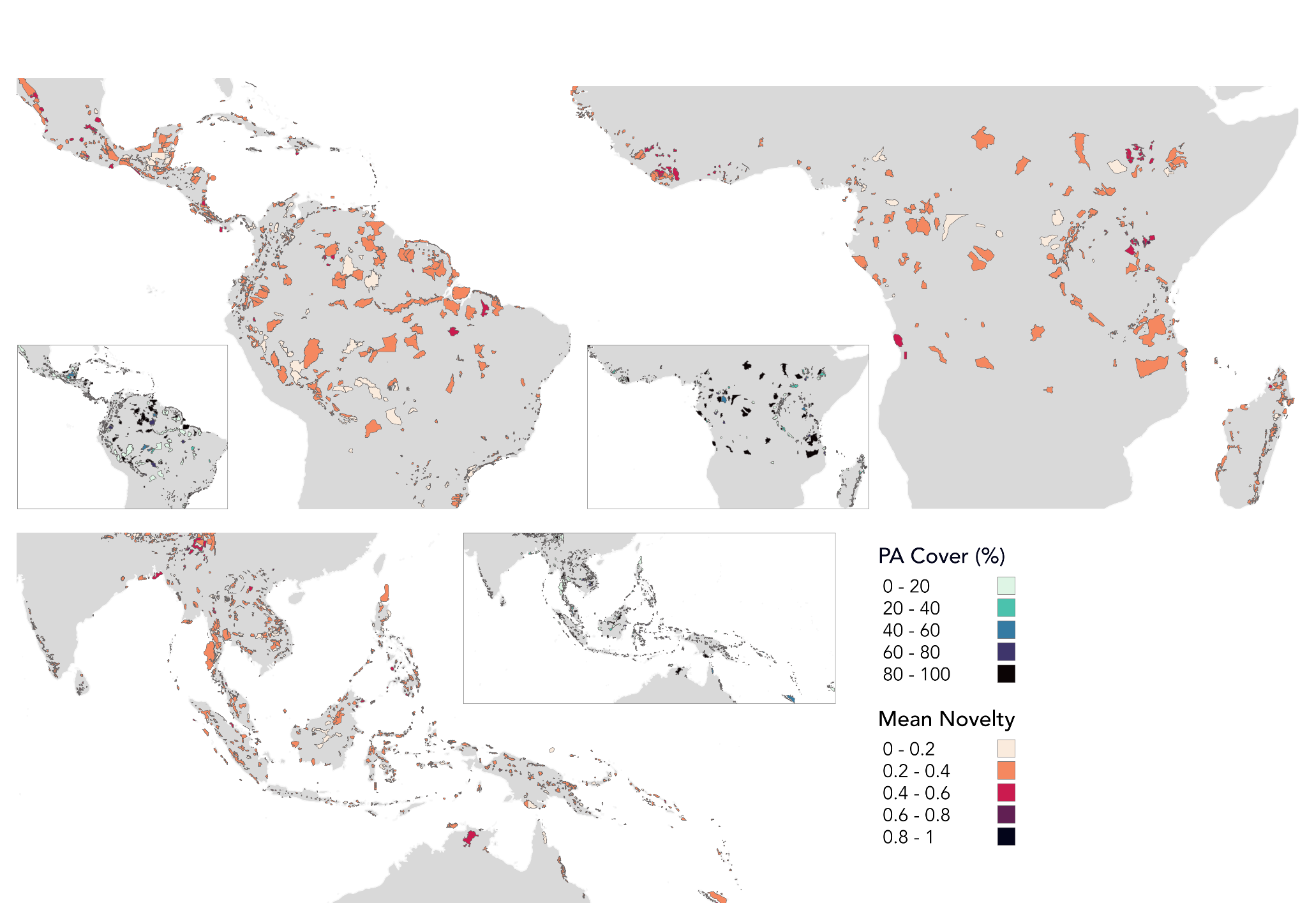

**Figure S2.2 |** Mean fractional novelty in recent temperature **isothermality** (2005 to 2019) compared to the historic baseline (1990 to 2004) across KBAs containing tropical forests in Latin America (n = 867), Africa (n = 395) and across Asia and Oceania (n = 1,262): 0-1, where 1 indicates entirely novel mean annual temperature regimes in 2005 - 2019. Individual inset maps show protected area coverage (%) for each KBA.

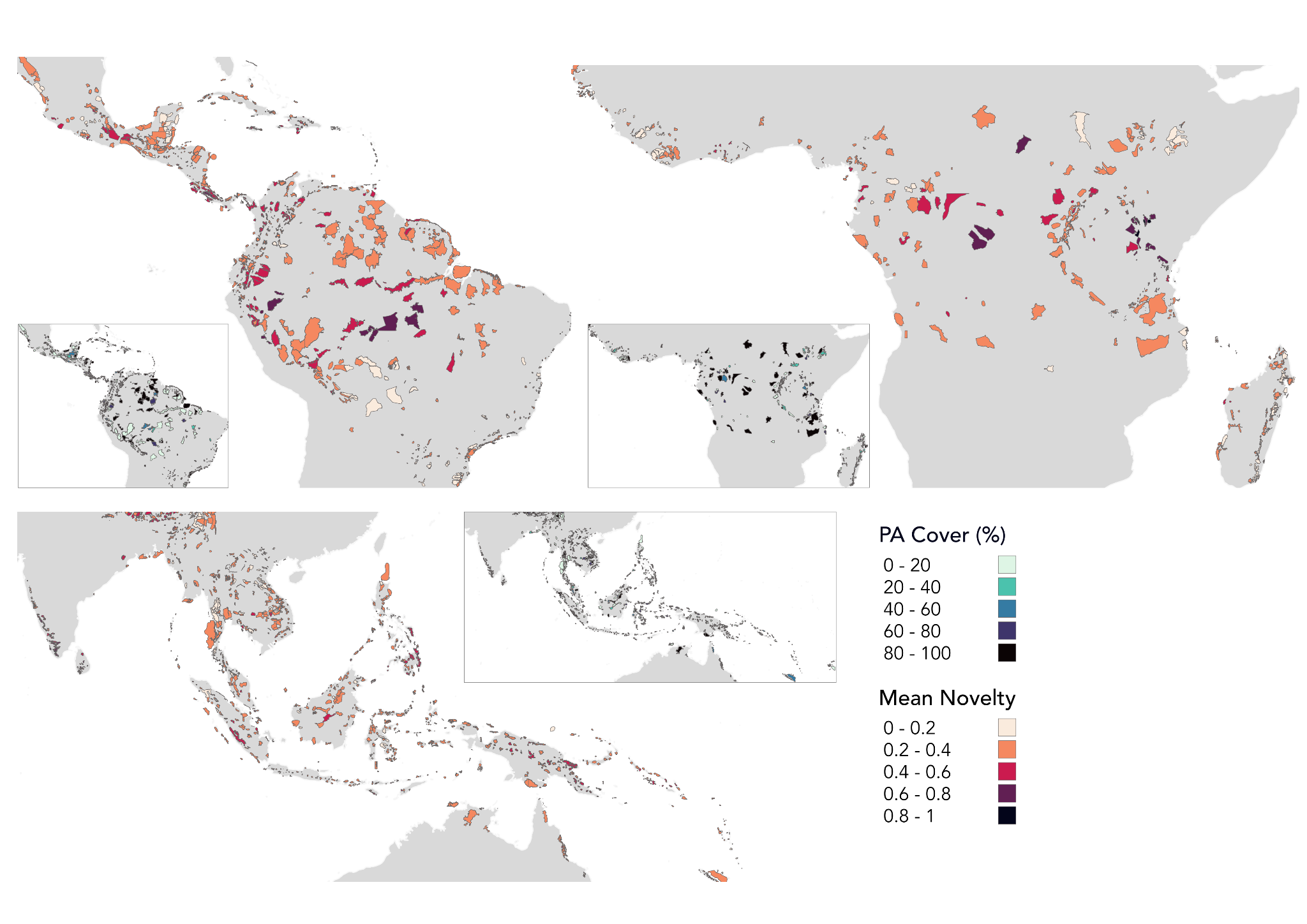

**Figure S2.3 |** Mean fractional novelty in recent **temperature seasonality** (2005 to 2019) compared to the historic baseline (1990 to 2004) across KBAs containing tropical forests in Latin America (n = 867), Africa (n = 395) and across Asia and Oceania (n = 1,262): 0-1, where 1 indicates entirely novel mean annual temperature regimes in 2005 - 2019. Individual inset maps show protected area coverage (%) for each KBA.

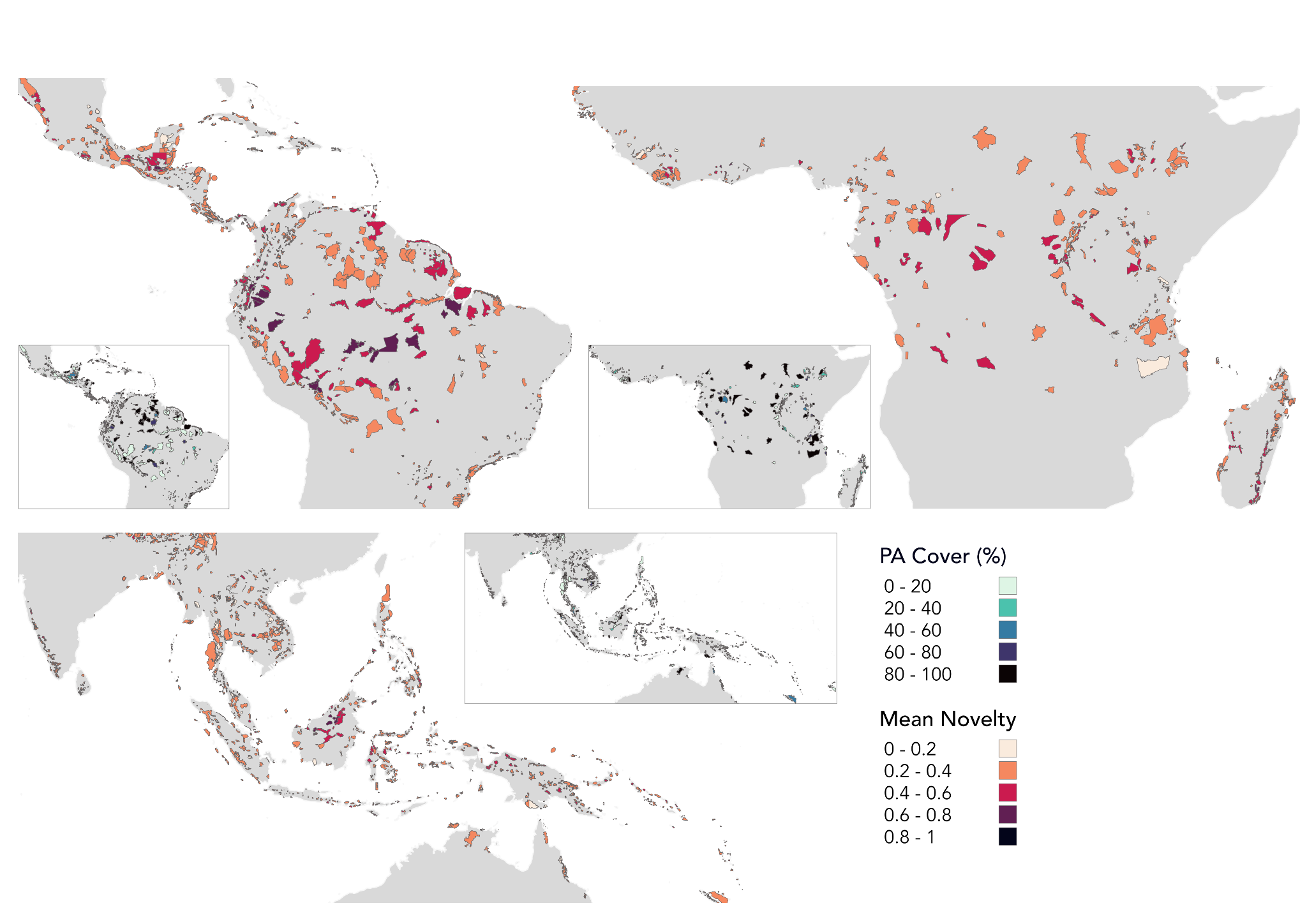

**Figure S2.4 |** Mean fractional novelty in recent **maximum temperature of the warmest month** (2005 to 2019) compared to the historic baseline (1990 to 2004) across KBAs containing tropical forests in Latin America (n = 867), Africa (n = 395) and across Asia and Oceania (n = 1,262): 0-1, where 1 indicates entirely novel mean annual temperature regimes in 2005 - 2019. Individual inset maps show protected area coverage (%) for each KBA.

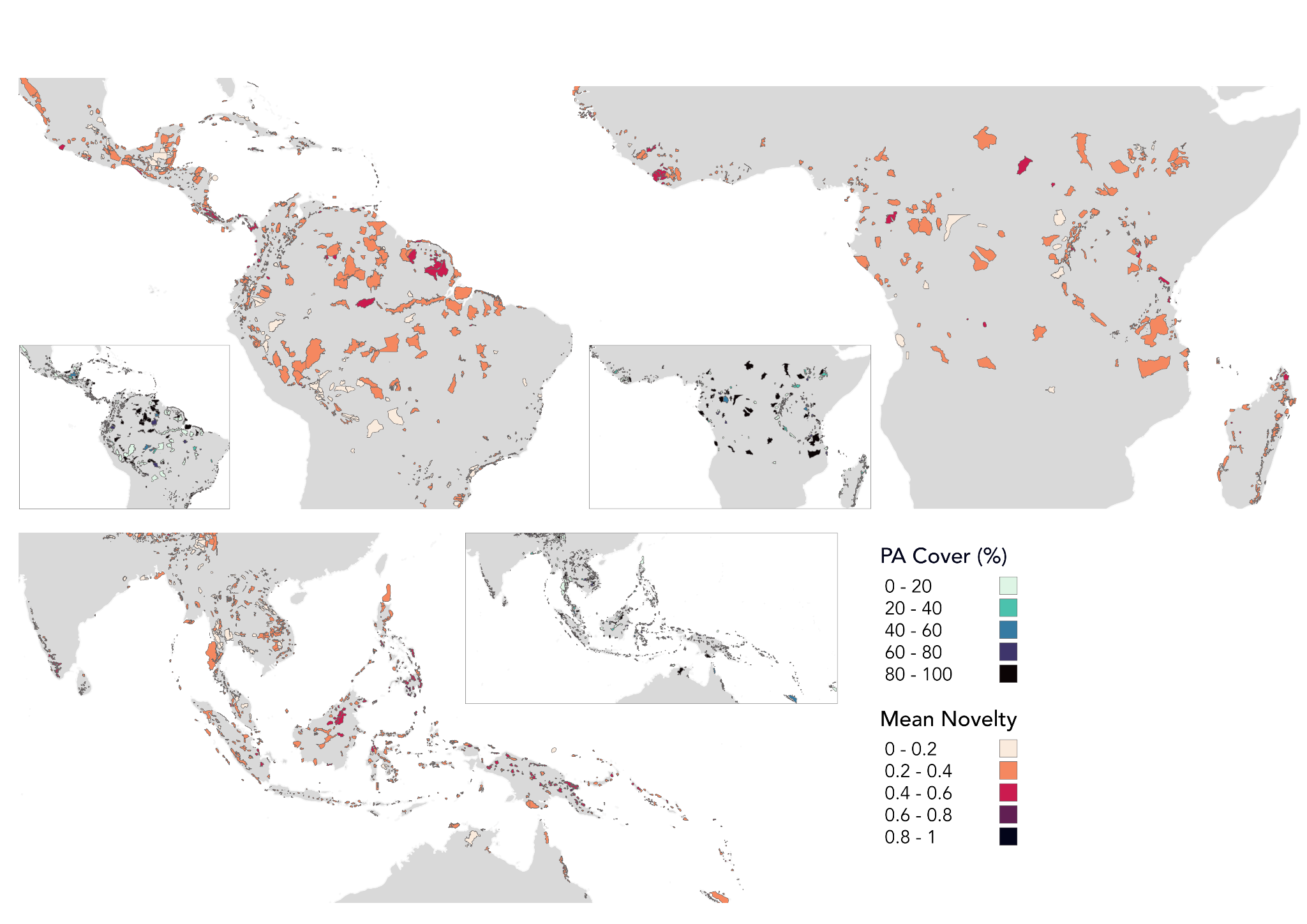

**Figure S2.5 |** Mean fractional novelty in recent **minimum temperature of the coldest month** (2005 to 2019) compared to the historic baseline (1990 to 2004) across KBAs containing tropical forests in Latin America (n = 867), Africa (n = 395) and across Asia and Oceania (n = 1,262): 0-1, where 1 indicates entirely novel mean annual temperature regimes in 2005 - 2019. Individual inset maps show protected area coverage (%) for each KBA.

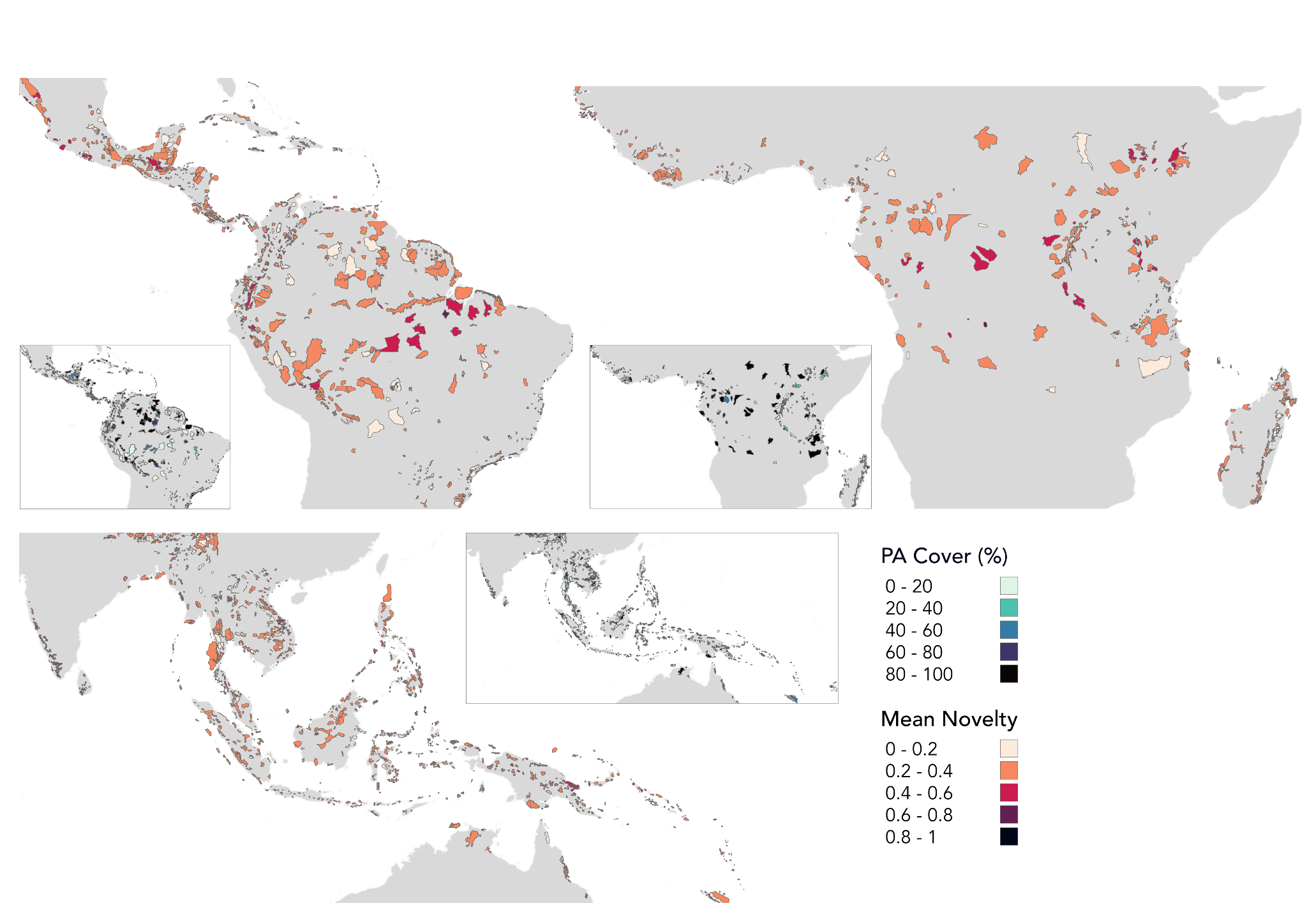

**Figure S2.6 |** Mean fractional novelty in recent **annual temperature range** (2005 to 2019) compared to the historic baseline (1990 to 2004) across KBAs containing tropical forests in Latin America (n = 867), Africa (n = 395) and across Asia and Oceania (n = 1,262): 0-1, where 1 indicates entirely novel mean annual temperature regimes in 2005 - 2019. Individual inset maps show protected area coverage (%) for each KBA.

**A**
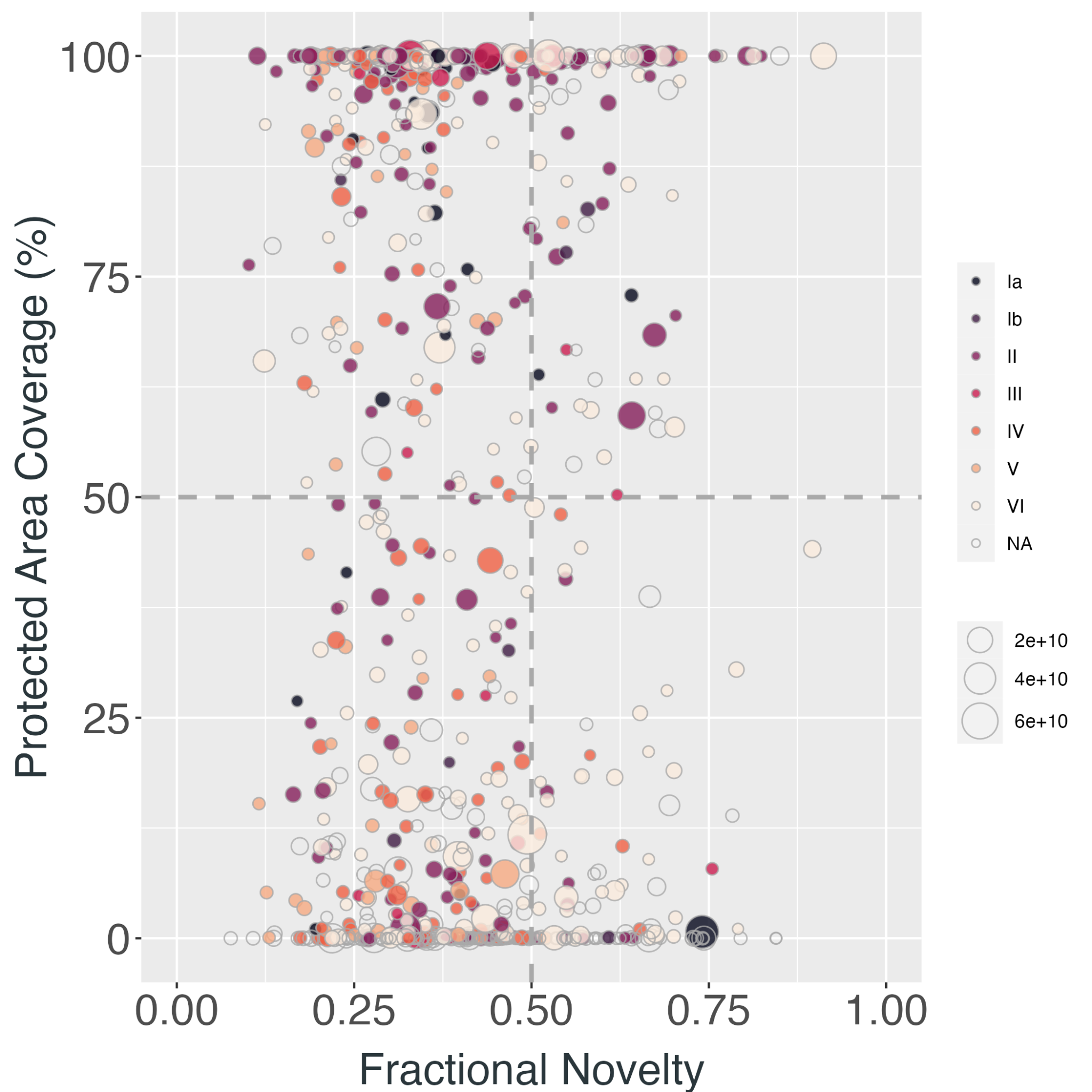
 **B**
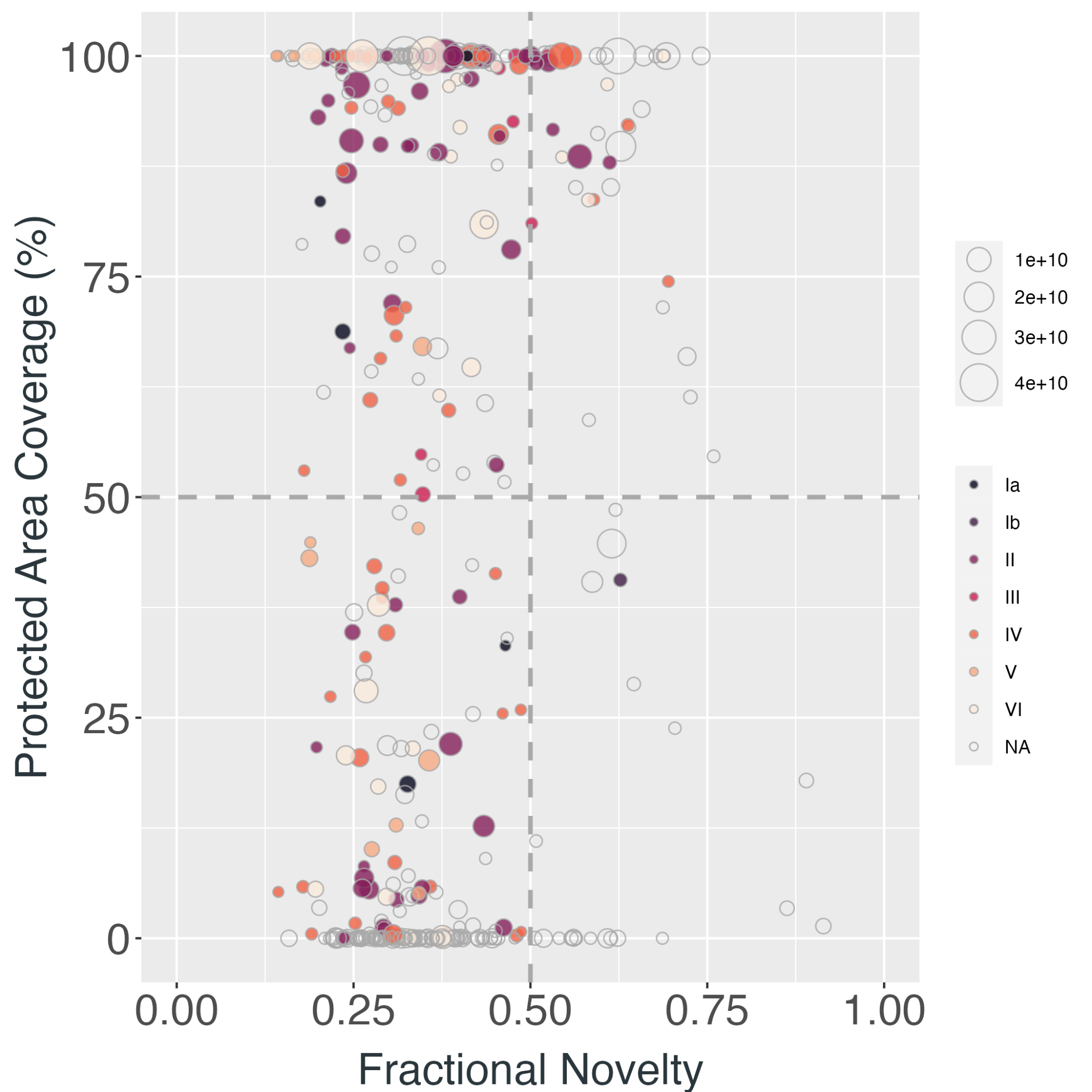
 **C**
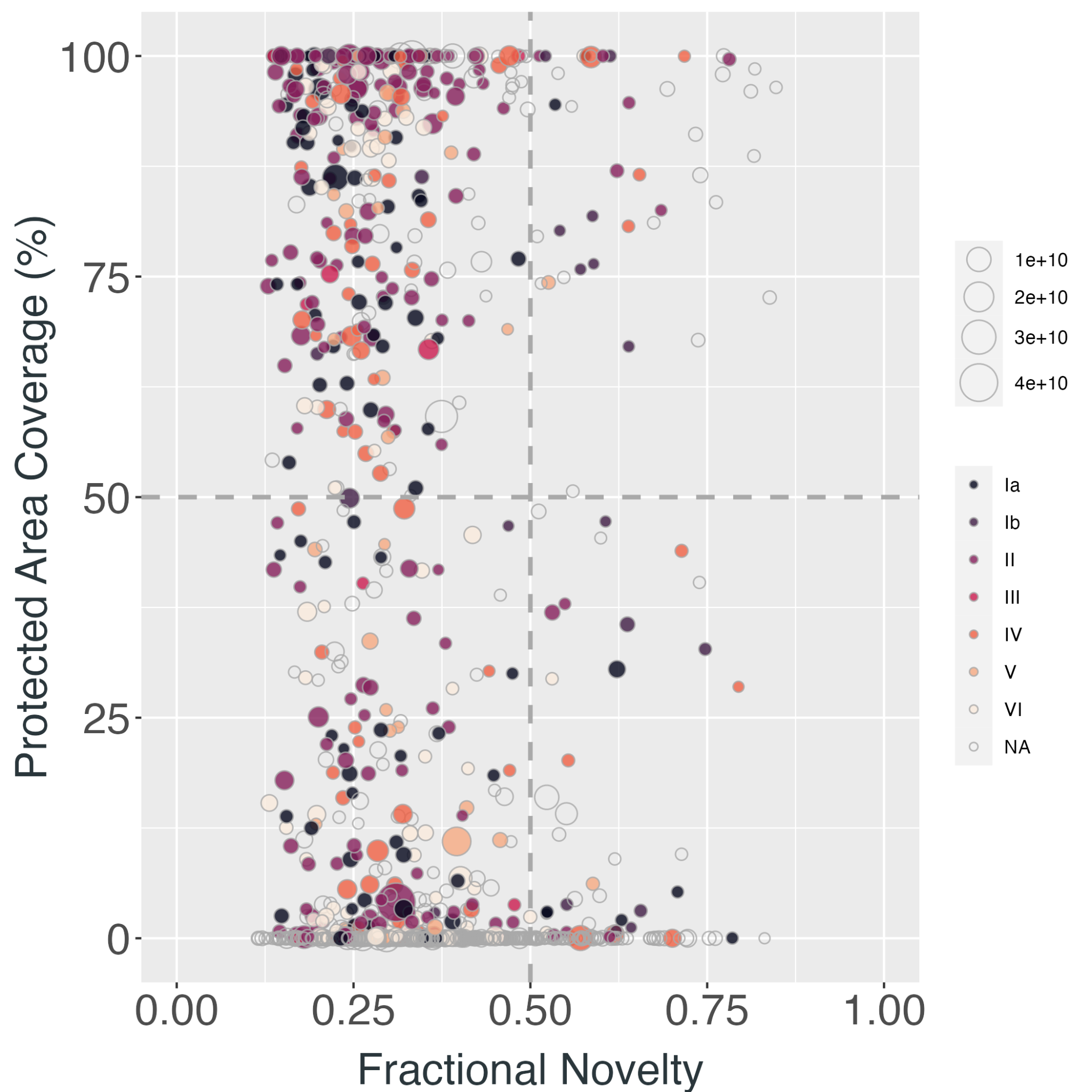

**Figure S3.1 |** Each point represents a Key Biodiversity Area (KBA) containing tropical forest in (A) Latin America, (B) Africa, (C) Asia and Australia and the relationship of the mean novelty in recent **mean diurnal temperature ranges** experienced by said Key Biodiversity Area and the formal protected area coverage (%) of the KBA. Point colours correspond to the IUCN category of the protected area and the size of each point corresponds to the geographical size of the KBA (m^2^).

**A**
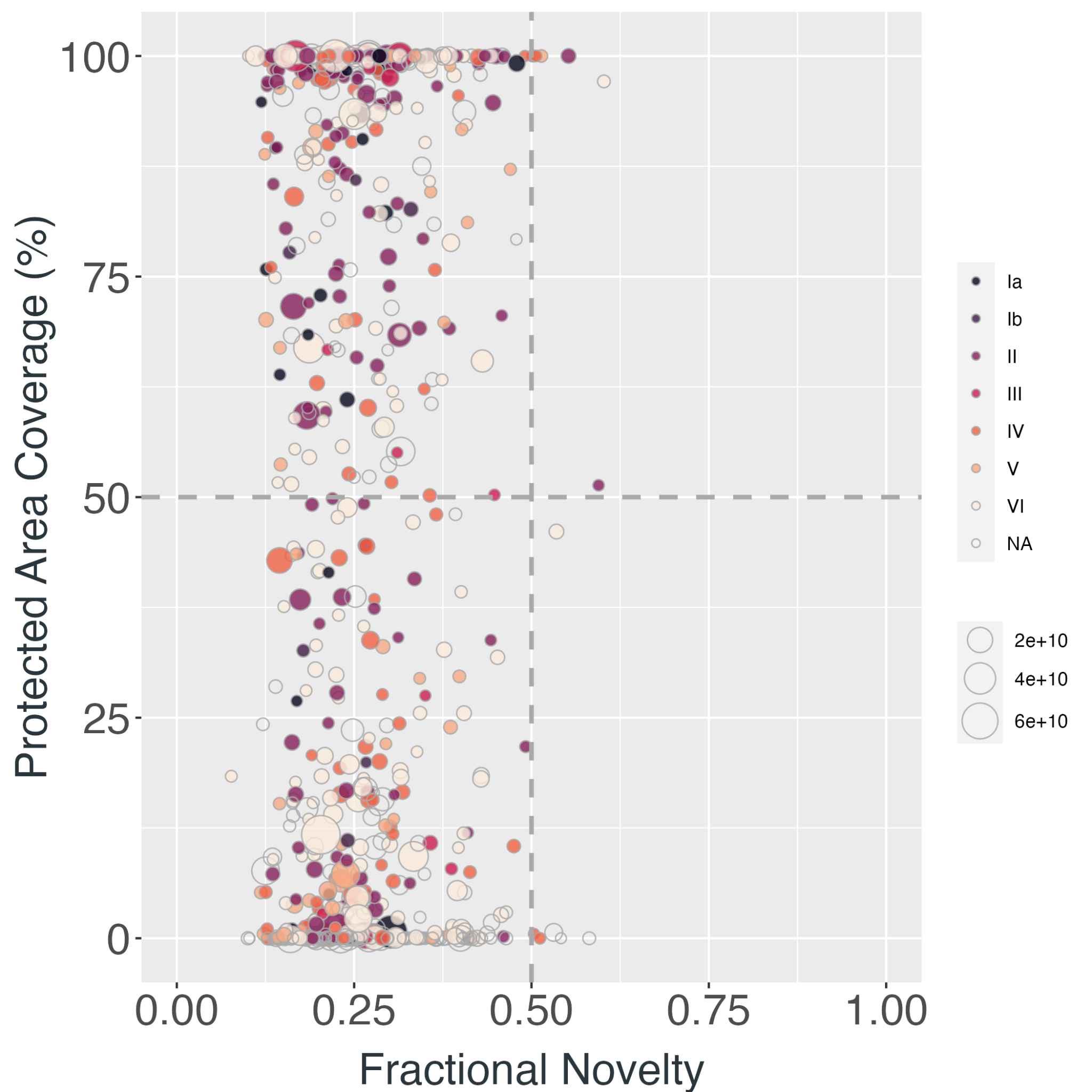
 **B**
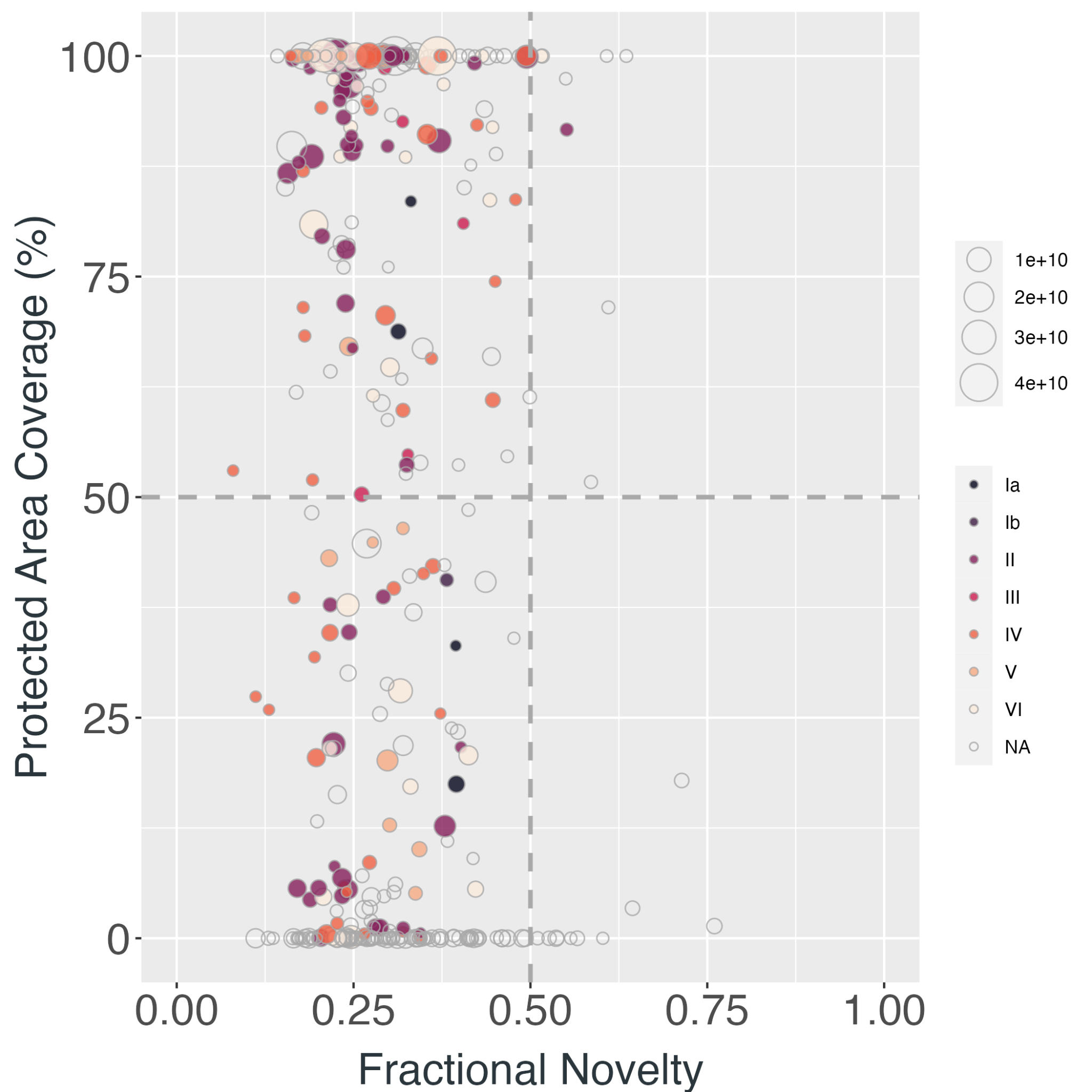
 **C**
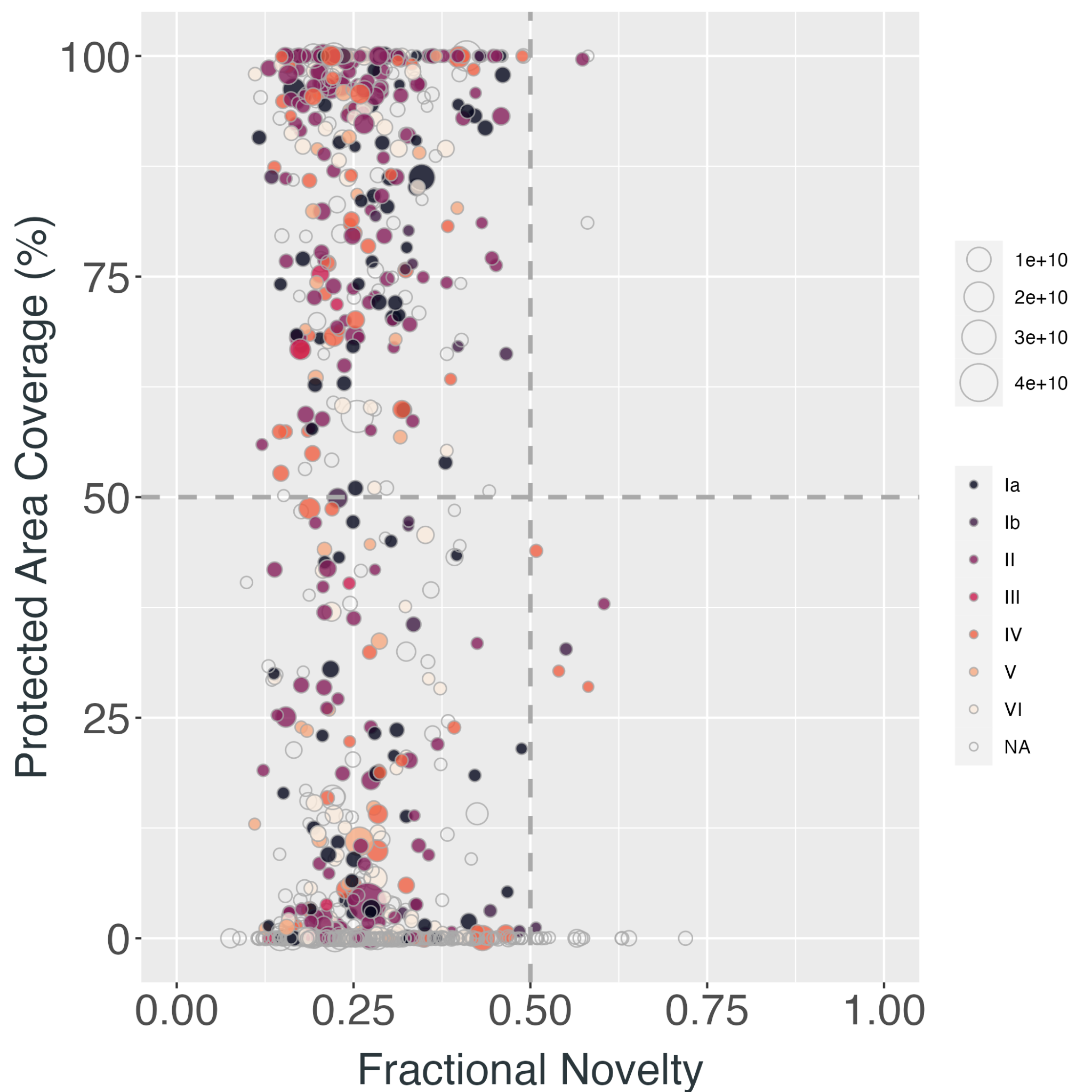

**Figure S3.2 |** Each point represents a Key Biodiversity Area (KBA) containing tropical forest in (A) Latin America, (B) Africa, (C) Asia and Australia and the relationship of the mean novelty in recent **isothermality** experienced by said Key Biodiversity Area and the formal protected area coverage (%) of the KBA. Point colours correspond to the IUCN category of the protected area and the size of each point corresponds to the geographical size of the KBA (m^2^).

**A**
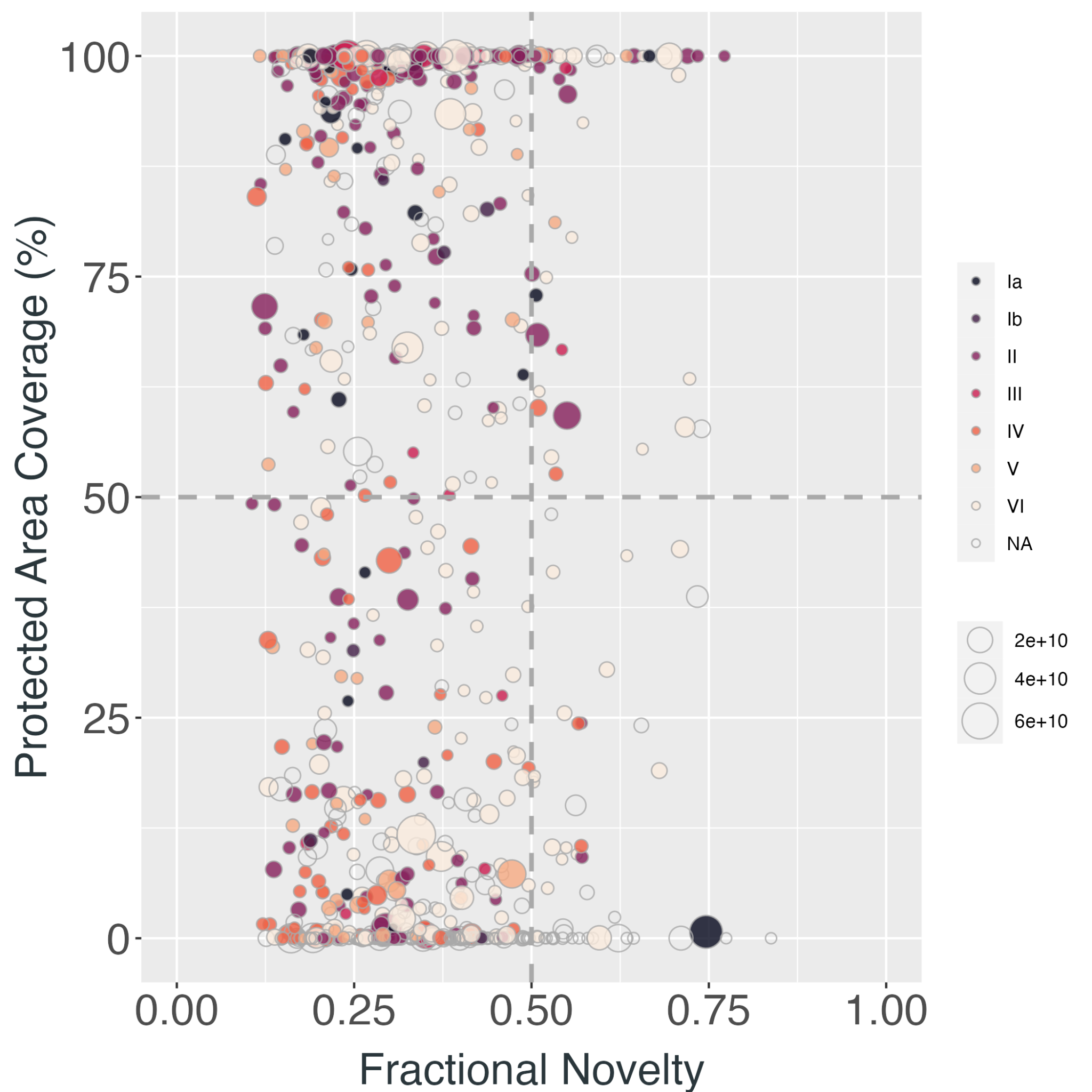
 **B**
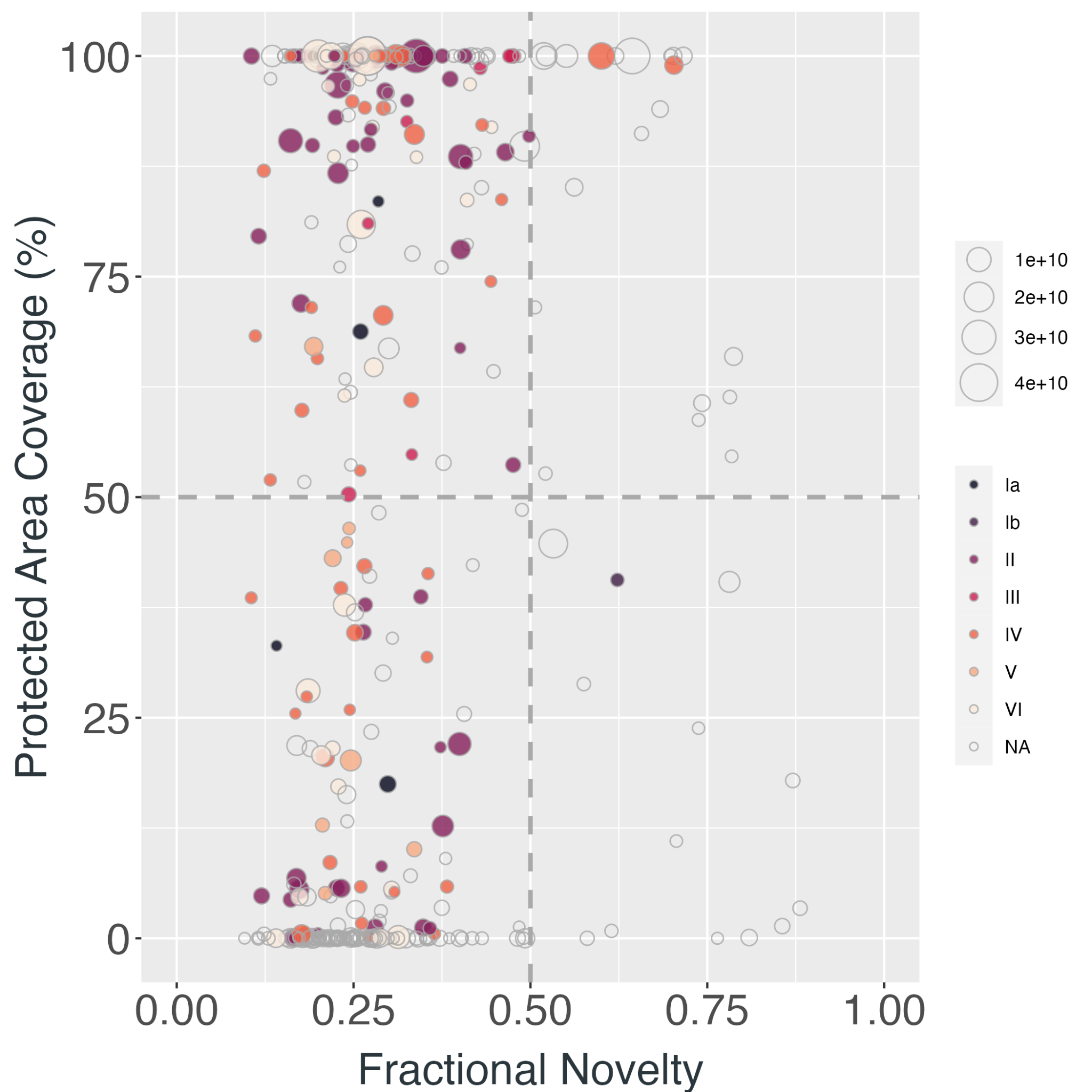
 **C**
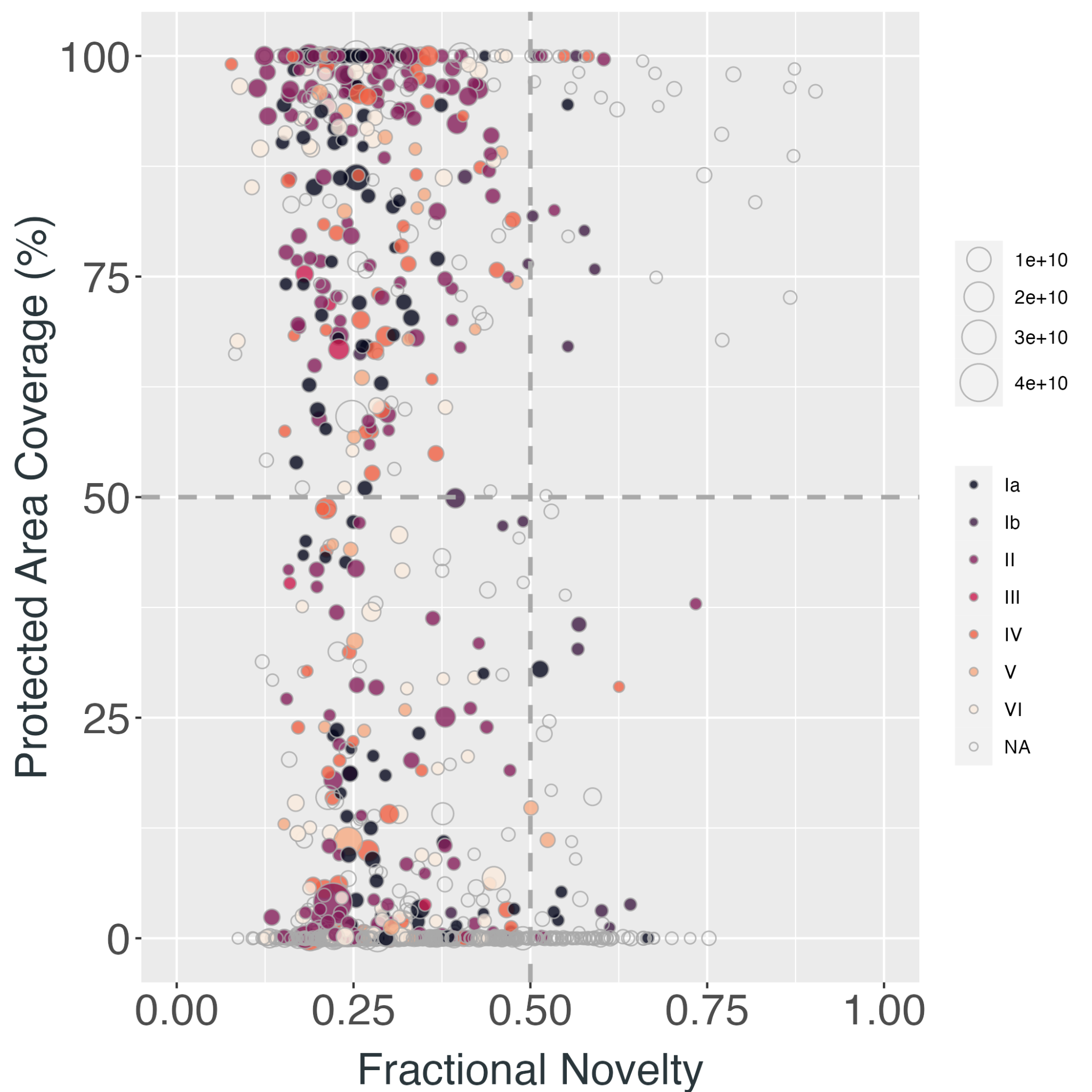

**Figure S3.3 |** Each point represents a Key Biodiversity Area (KBA) containing tropical forest in (A) Latin America, (B) Africa, (C) Asia and Australia and the relationship of the mean novelty in recent **temperature seasonality** experienced by said Key Biodiversity Area and the formal protected area coverage (%) of the KBA. Point colours correspond to the IUCN category of the protected area and the size of each point corresponds to the geographical size of the KBA (m^2^).

**A**
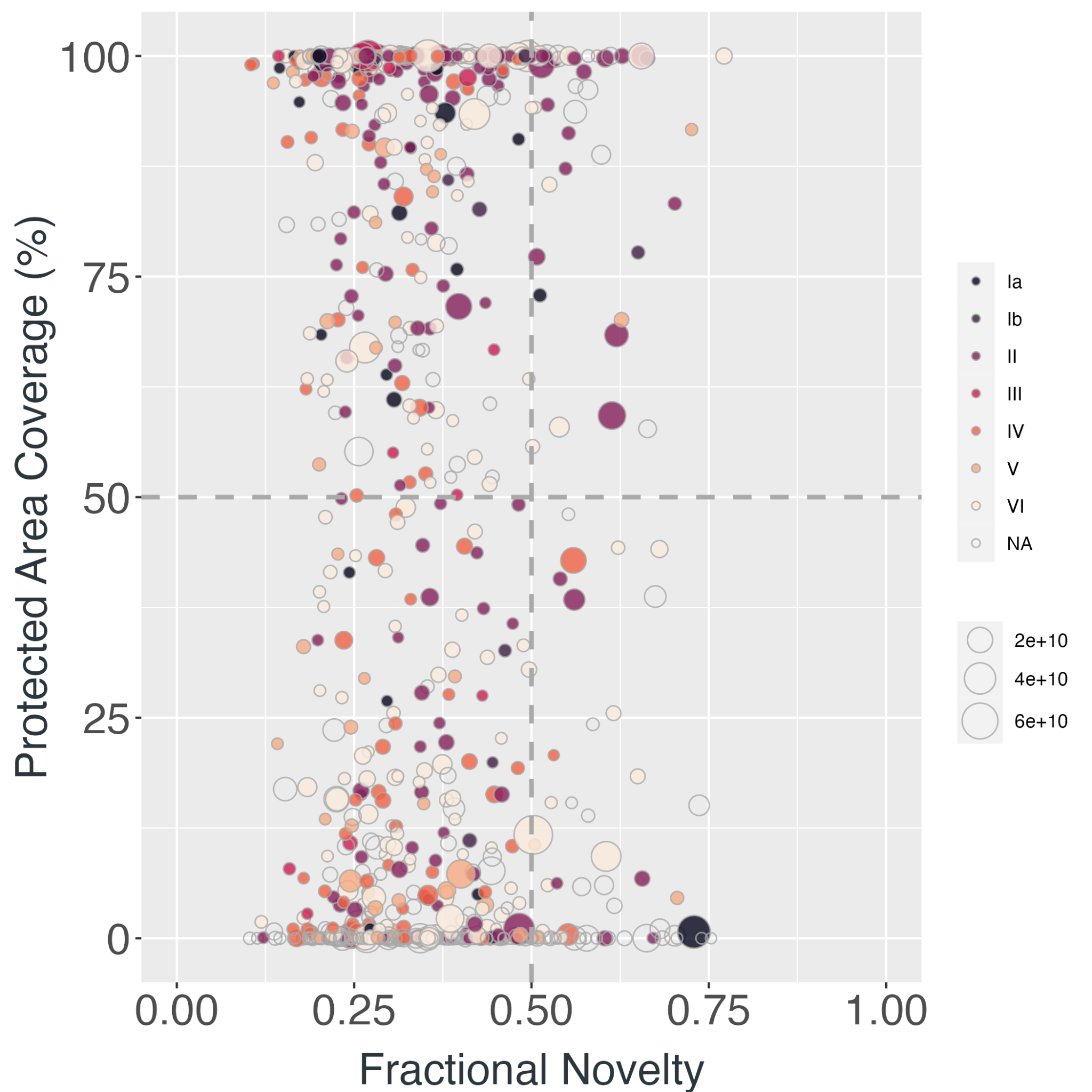
 **B**
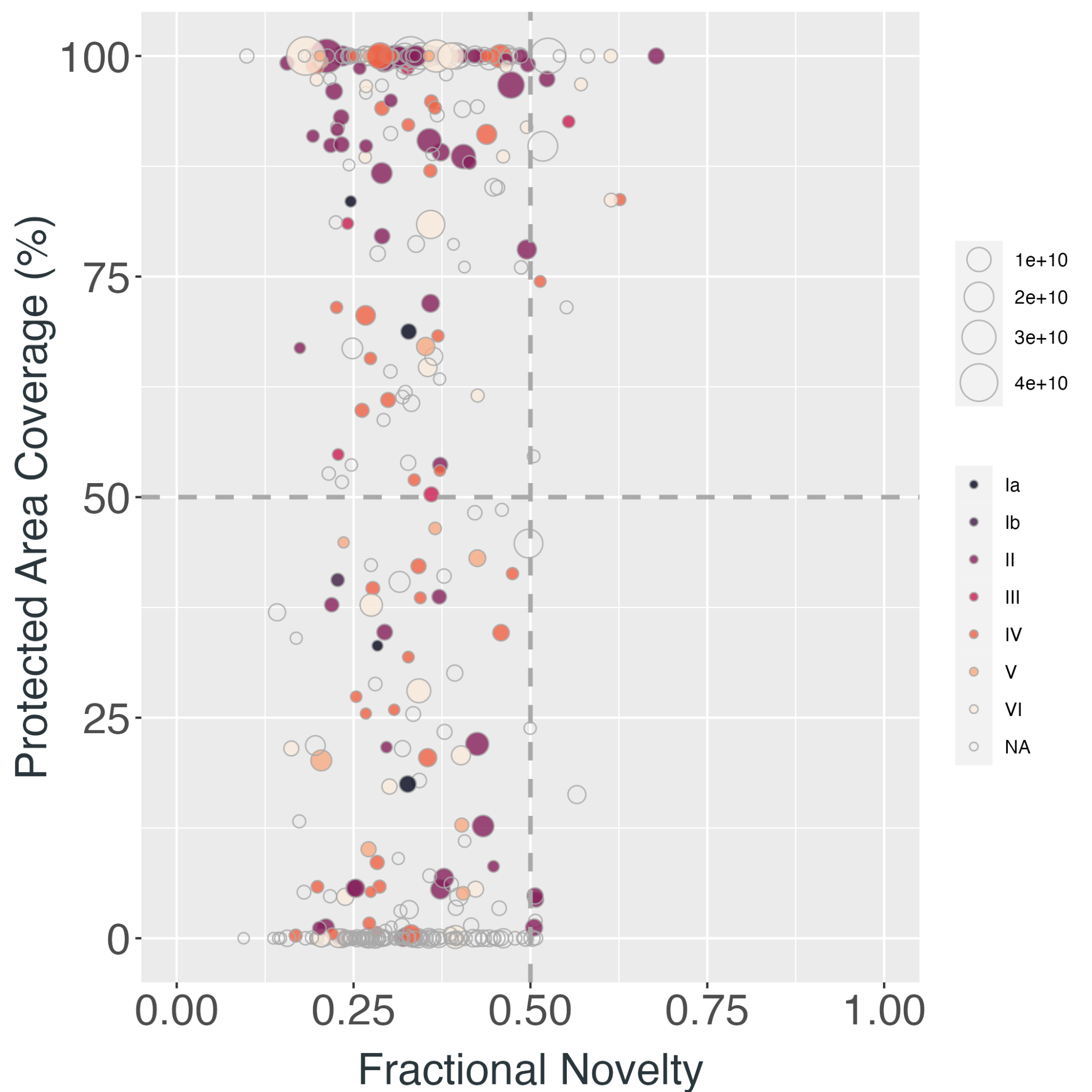
 **C**
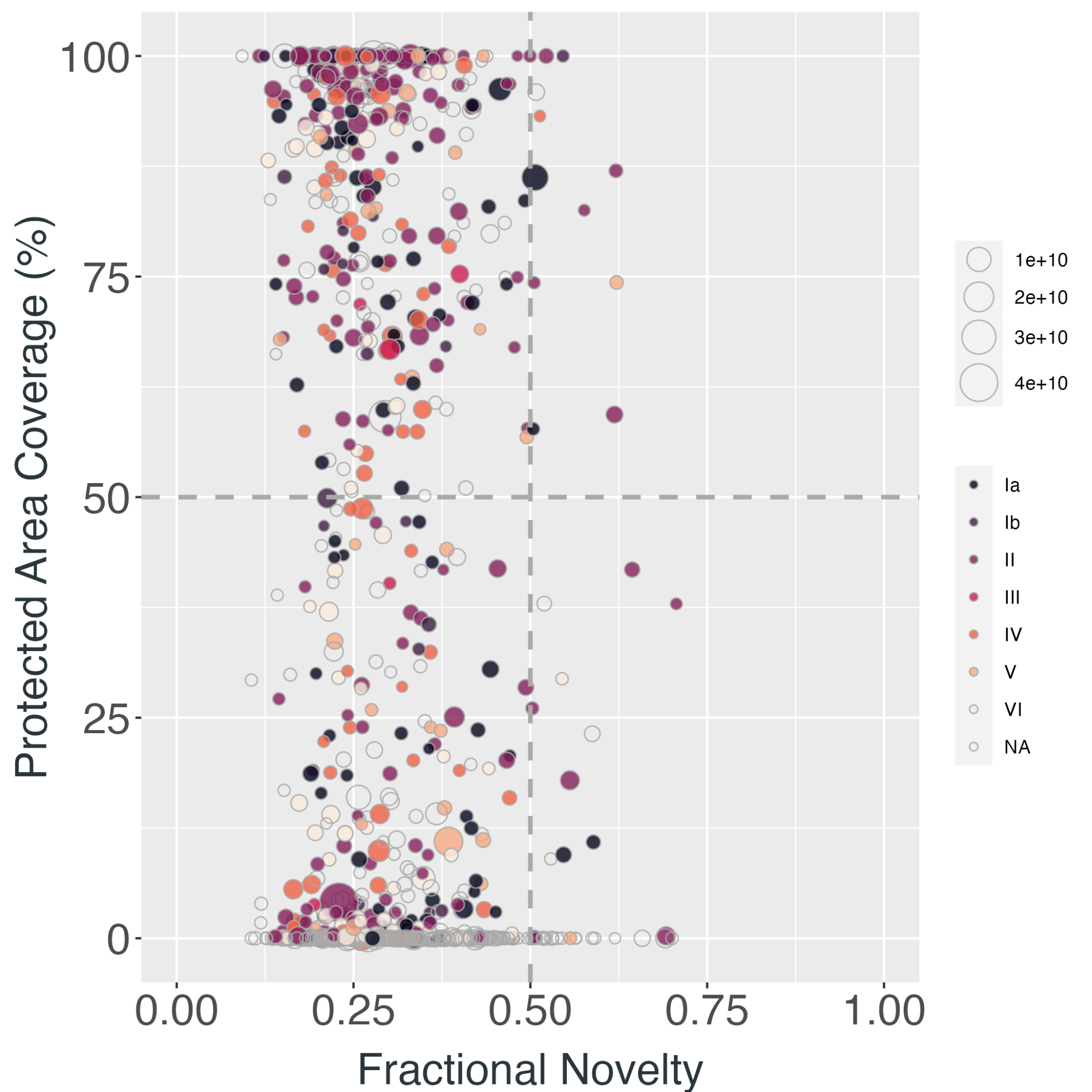

**Figure S3.4 |** Each point represents a Key Biodiversity Area (KBA) containing tropical forest in (A) Latin America, (B) Africa, (C) Asia and Australia and the relationship of the mean novelty in recent **maximum temperature of the warmest month** experienced by said Key Biodiversity Area and the formal protected area coverage (%) of the KBA. Point colours correspond to the IUCN category of the protected area and the size of each point corresponds to the geographical size of the KBA (m^2^).

**A**
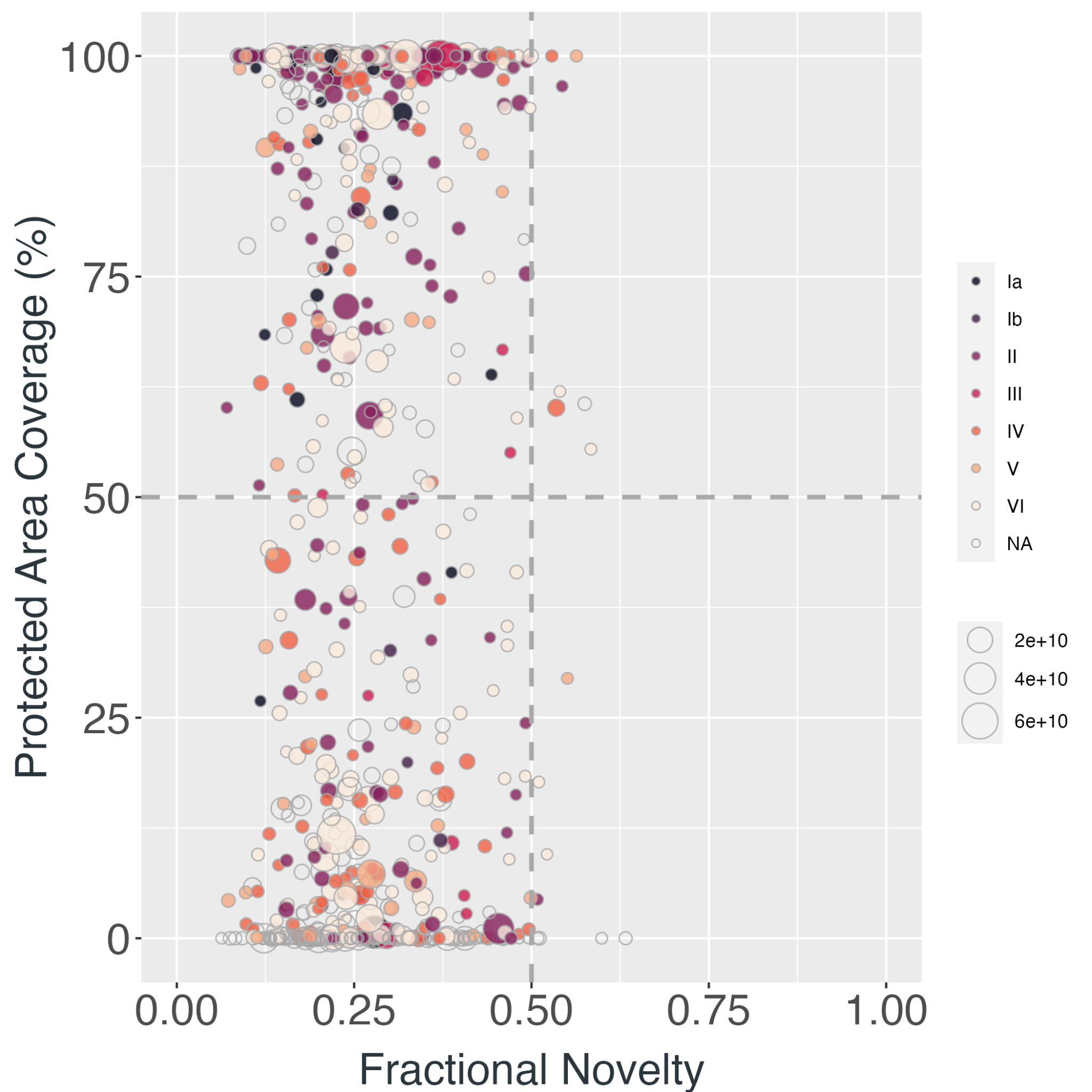
 **B**
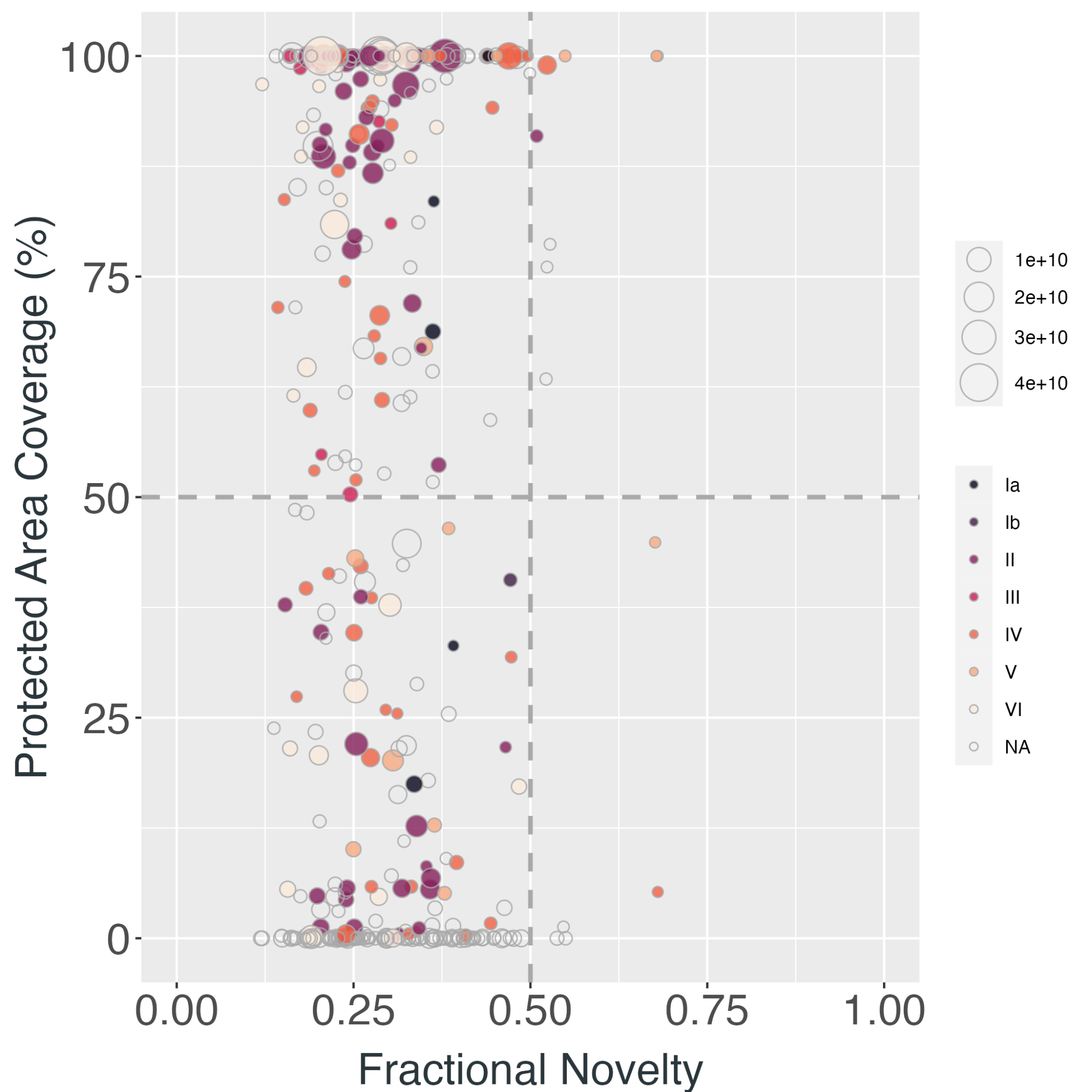
 **C**
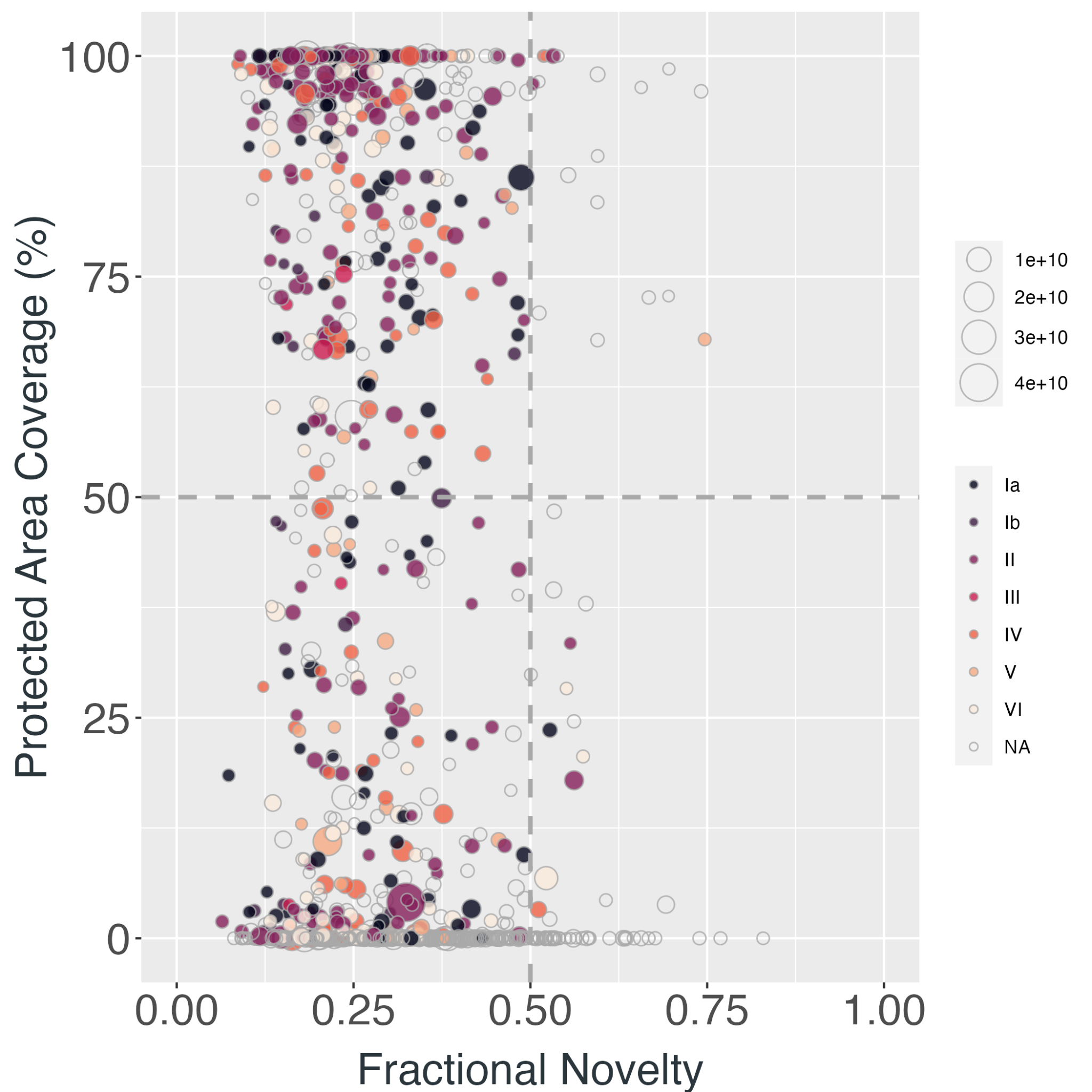

**Figure S3.5 |** Each point represents a Key Biodiversity Area (KBA) containing tropical forest in (A) Latin America, (B) Africa, (C) Asia and Australia and the relationship of the mean novelty in recent **minimum temperature of the coldest month** experienced by said Key Biodiversity Area and the formal protected area coverage (%) of the KBA. Point colours correspond to the IUCN category of the protected area and the size of each point corresponds to the geographical size of the KBA (m^2^).

**A**
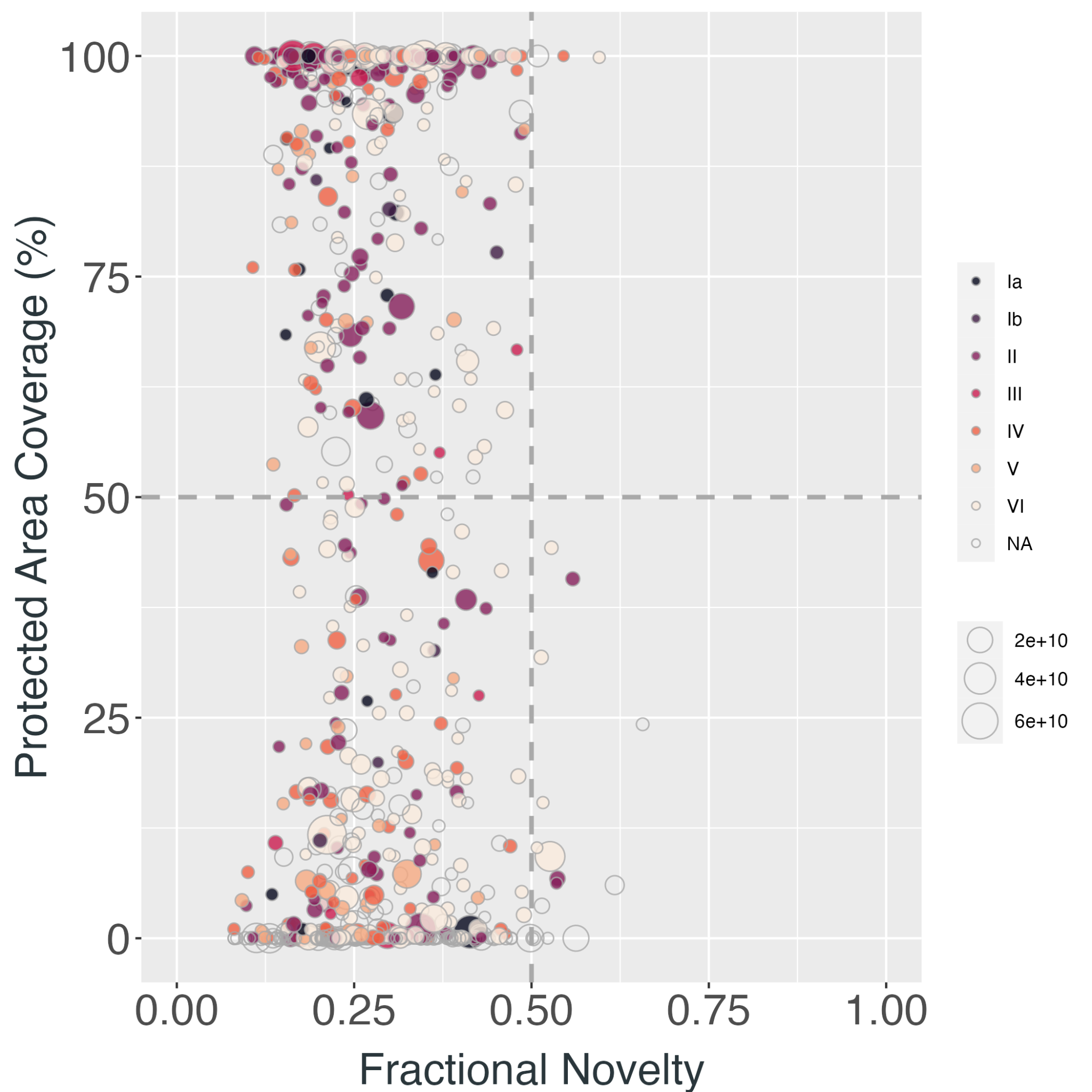
 **B**
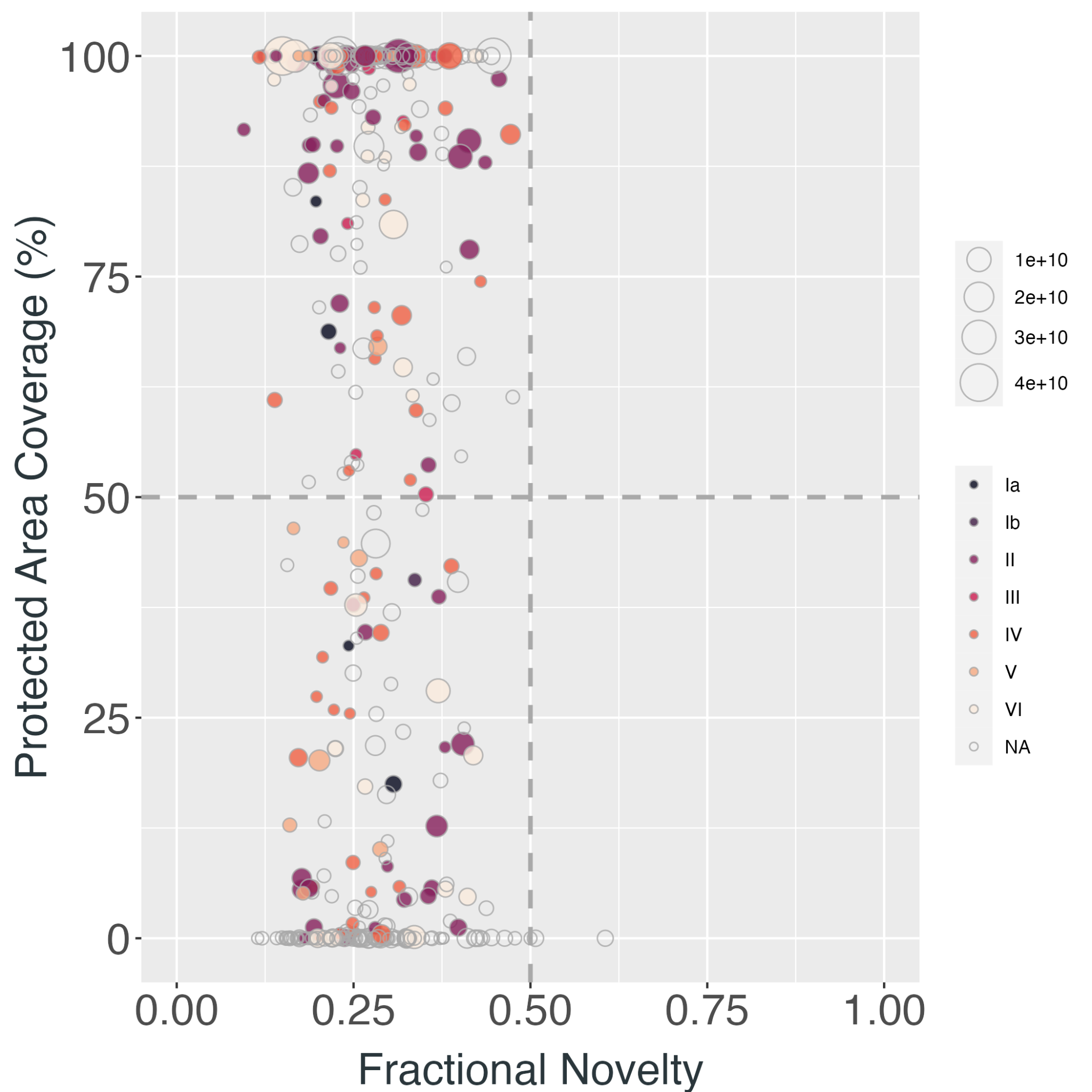
 **C**
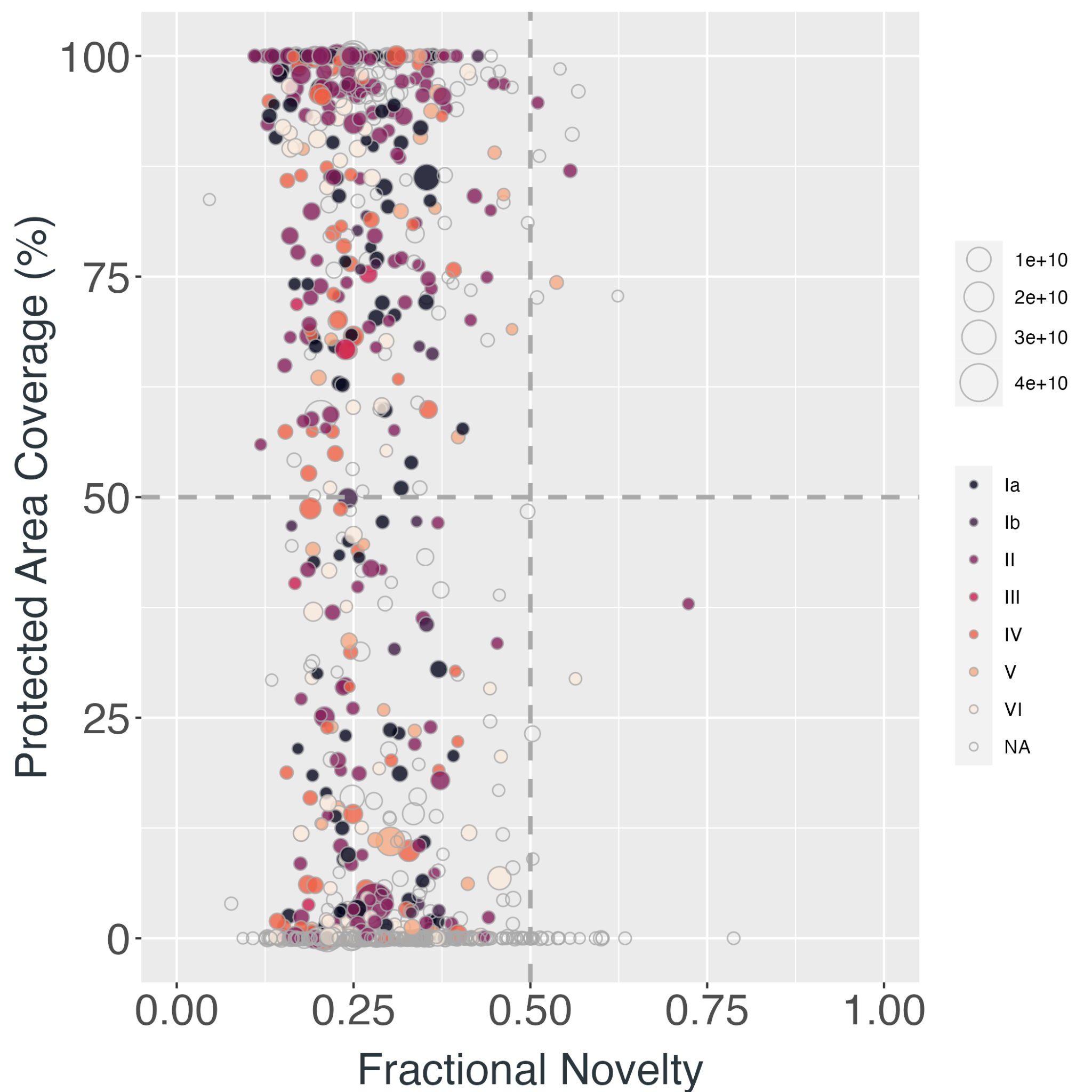

**Figure S3.6 |** Each point represents a Key Biodiversity Area (KBA) containing tropical forest in (A) Latin America, (B) Africa, (C) Asia and Australia and the relationship of the mean novelty in recent **annual temperature range** experienced by said Key Biodiversity Area and the formal protected area coverage (%) of the KBA. Point colours correspond to the IUCN category of the protected area and the size of each point corresponds to the geographical size of the KBA (m^2^).

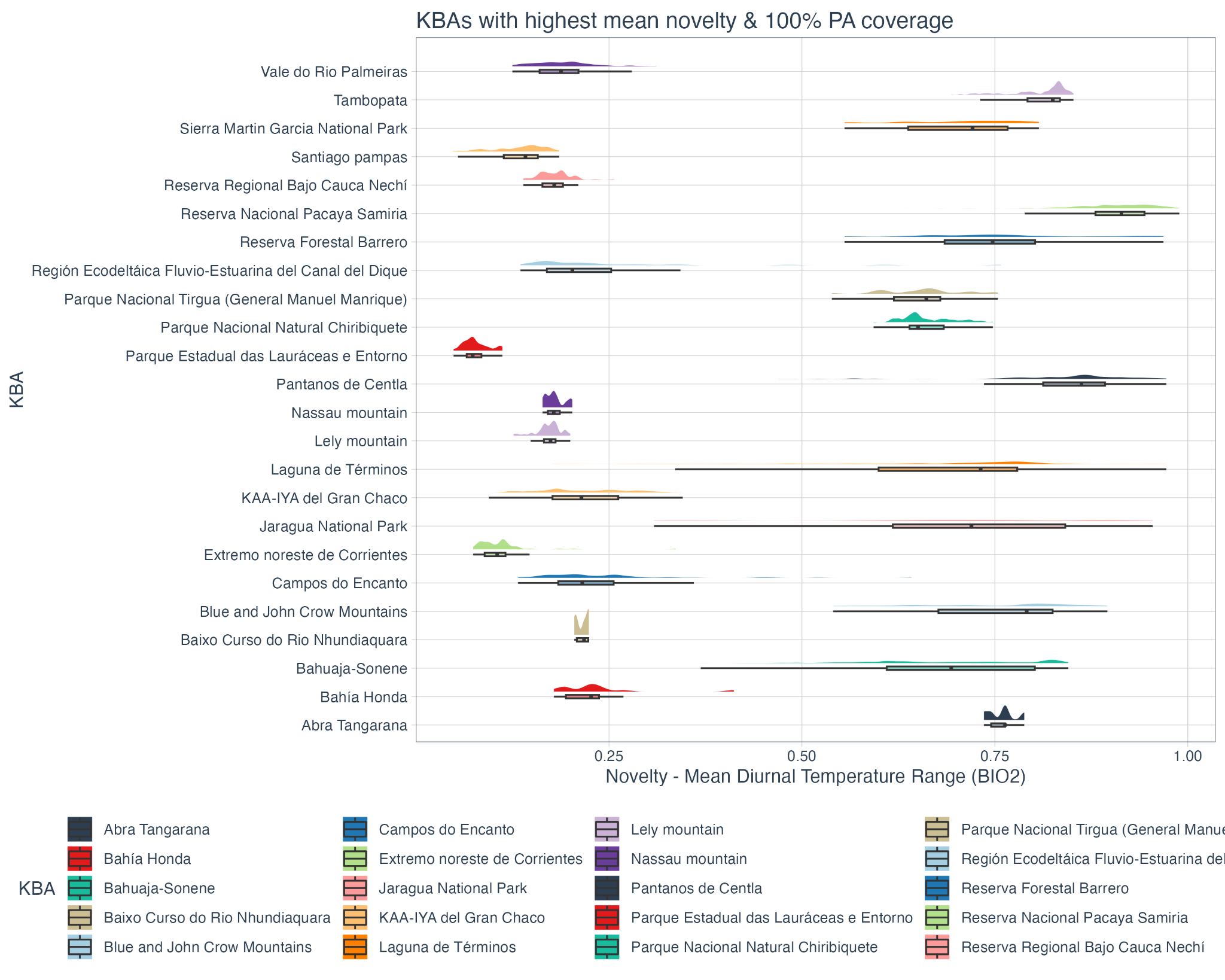

**Figure S4.1 |** Fractional novelty in recent mean diurnal temperature range (2005 to 2019) compared to the historic baseline (1990 to 2004) for the 12 KBAs in **Latin America** containing tropical forest with at least 99% protected area coverage and the highest recent mean fractional novelty in mean diurnal temperature range and the 12 KBAs in Latin America containing tropical forest with the lowest recent mean fractional novelty in mean diurnal temperature range and no protected area coverage. Novelty is measured between 0-1, where 1 indicates entirely novel mean diurnal temperature range regimes in 2005 - 2019.

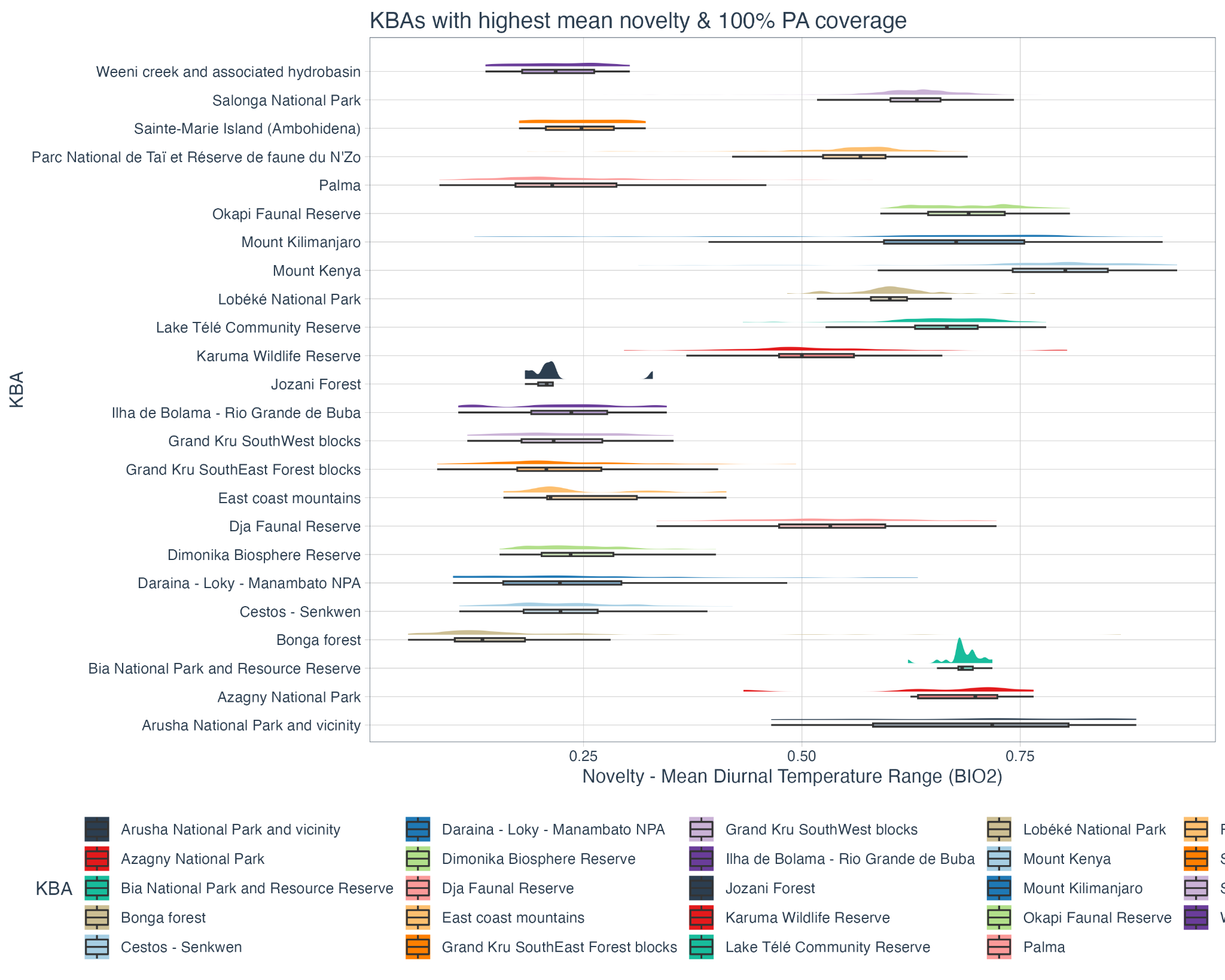

**Figure S4.2 |** Fractional novelty in recent mean diurnal temperature range (2005 to 2019) compared to the historic baseline (1990 to 2004) for the 12 KBAs in **Africa** containing tropical forest with at least 99% protected area coverage and the highest recent mean fractional novelty in mean diurnal temperature range and the 12 KBAs in Africa containing tropical forest with the lowest recent mean fractional novelty in mean diurnal temperature range and no protected area coverage. Novelty is measured between 0-1, where 1 indicates entirely novel mean diurnal temperature range regimes in 2005 - 2019.

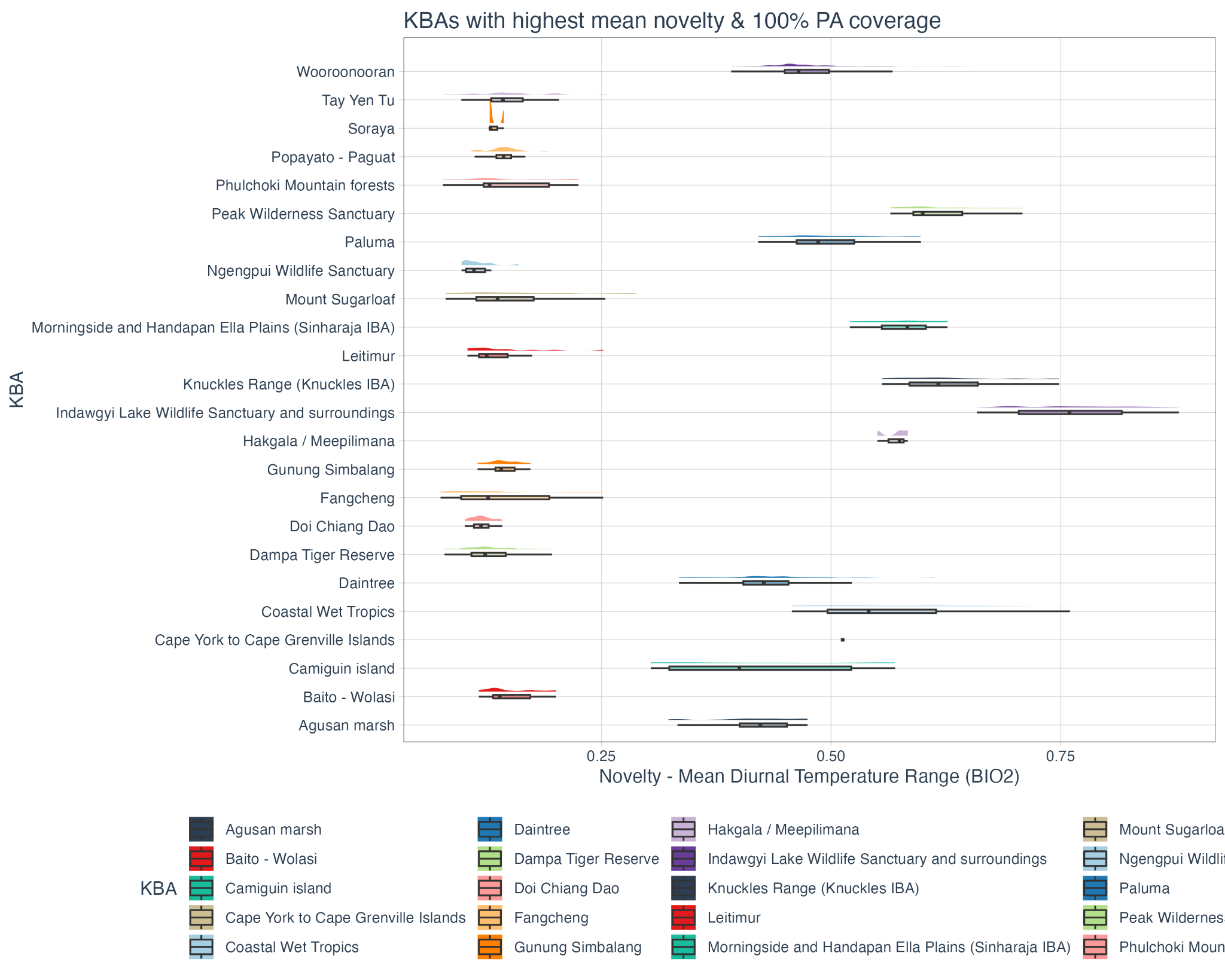

**Figure S4.3 |** Fractional novelty in recent mean diurnal temperature range (2005 to 2019) compared to the historic baseline (1990 to 2004) for the 12 KBAs in **Asia and Oceania** containing tropical forest with at least 99% protected area coverage and the highest recent mean fractional novelty in mean diurnal temperature range and the 12 KBAs in Asia and Oceania containing tropical forest with the lowest recent mean fractional novelty in mean diurnal temperature range and no protected area coverage. Novelty is measured between 0-1, where 1 indicates entirely novel mean diurnal temperature range regimes in 2005 - 2019.

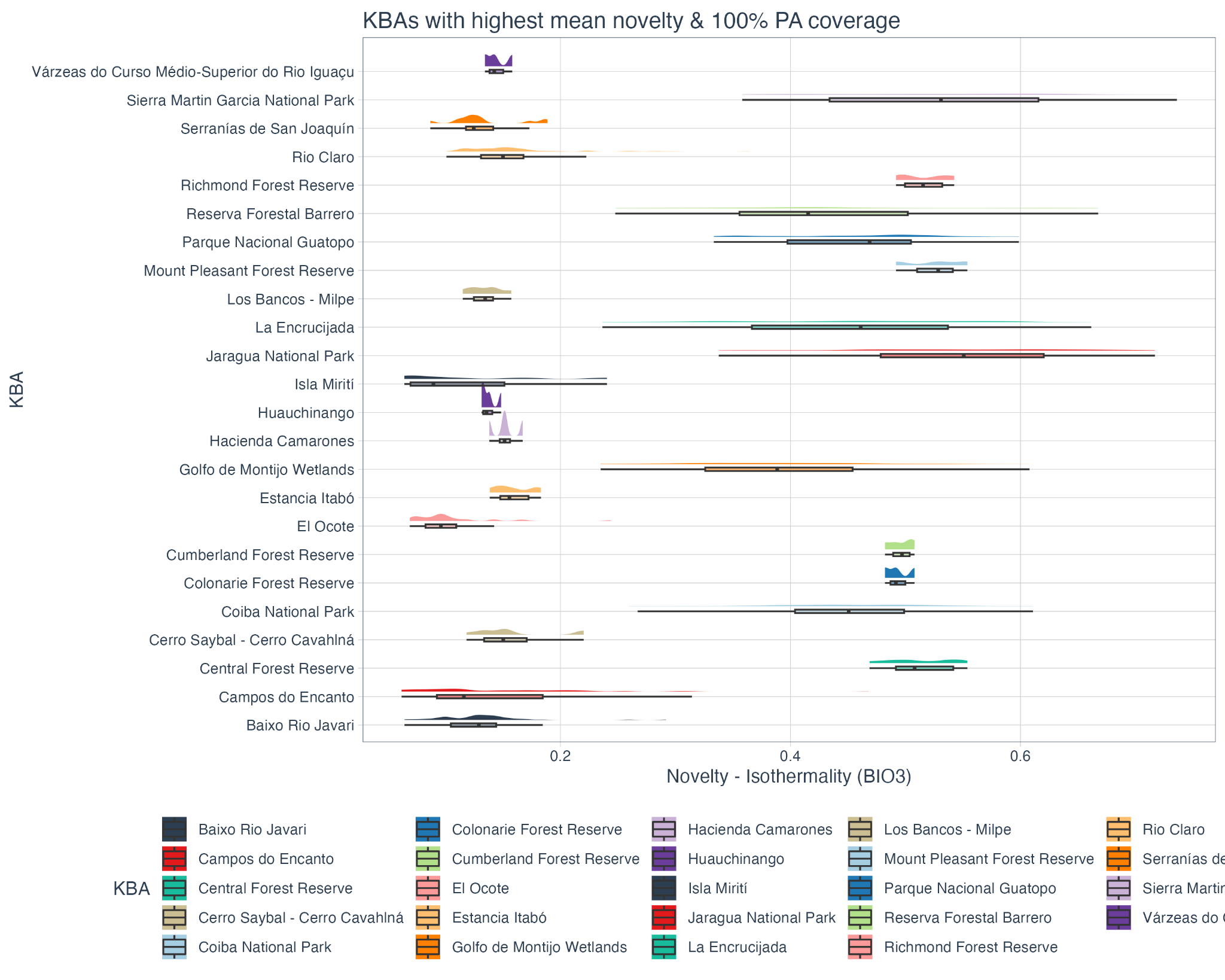

**Figure S4.4 |** Fractional novelty in recent isothermality (2005 to 2019) compared to the historic baseline (1990 to 2004) for the 12 KBAs in **Latin America** containing tropical forest with at least 99% protected area coverage and the highest recent mean fractional novelty in isothermality and the 12 KBAs in Latin America containing tropical forest with the lowest recent mean fractional novelty in isothermality and no protected area coverage. Novelty is measured between 0-1, where 1 indicates entirely novel isothermality regimes in 2005 - 2019.

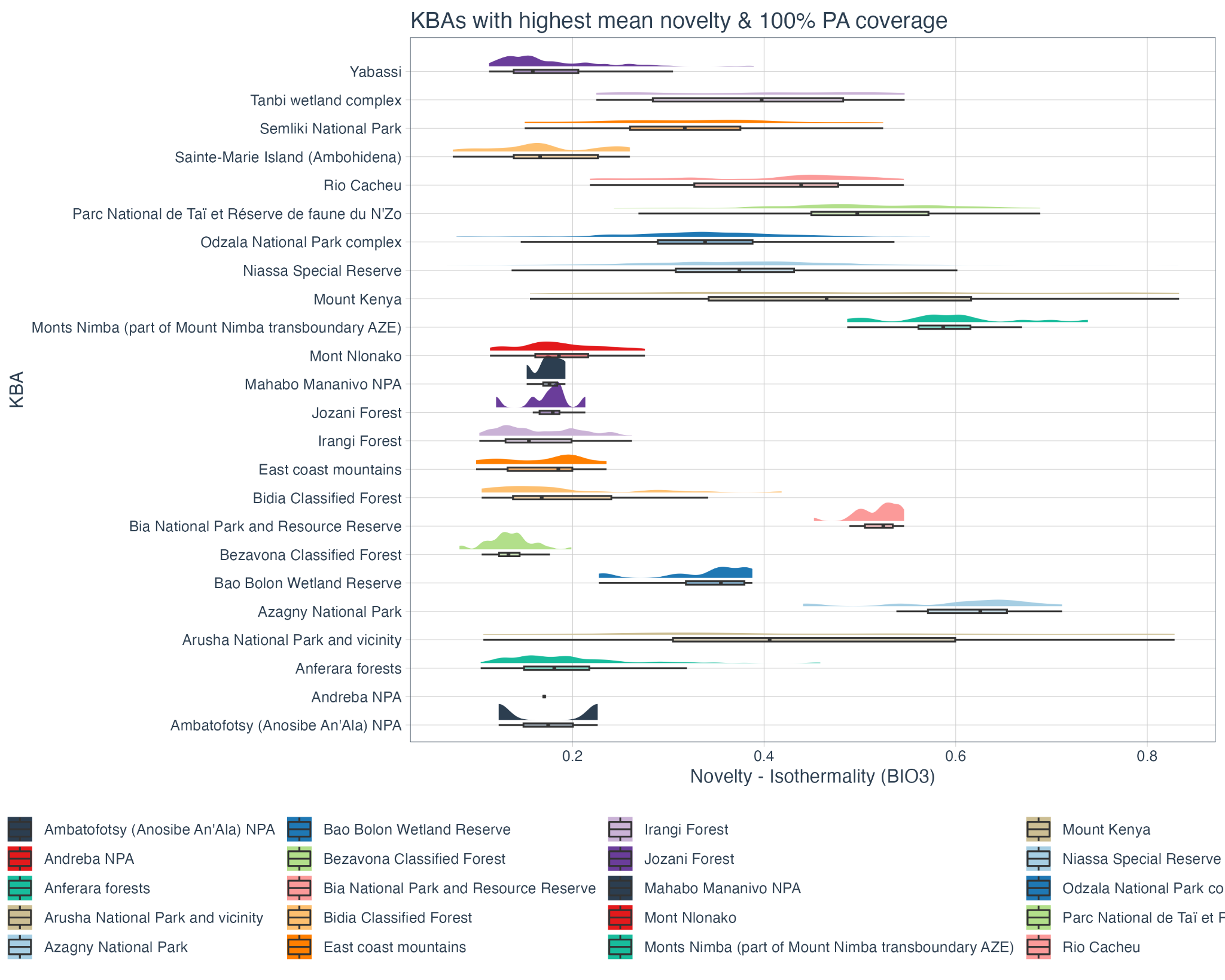

**Figure S4.5 |** Fractional novelty in recent isothermality (2005 to 2019) compared to the historic baseline (1990 to 2004) for the 12 KBAs in **Africa** containing tropical forest with at least 99% protected area coverage and the highest recent mean fractional novelty in isothermality and the 12 KBAs in Africa containing tropical forest with the lowest recent mean fractional novelty in isothermality and no protected area coverage. Novelty is measured between 0-1, where 1 indicates entirely novel isothermality regimes in 2005 - 2019.

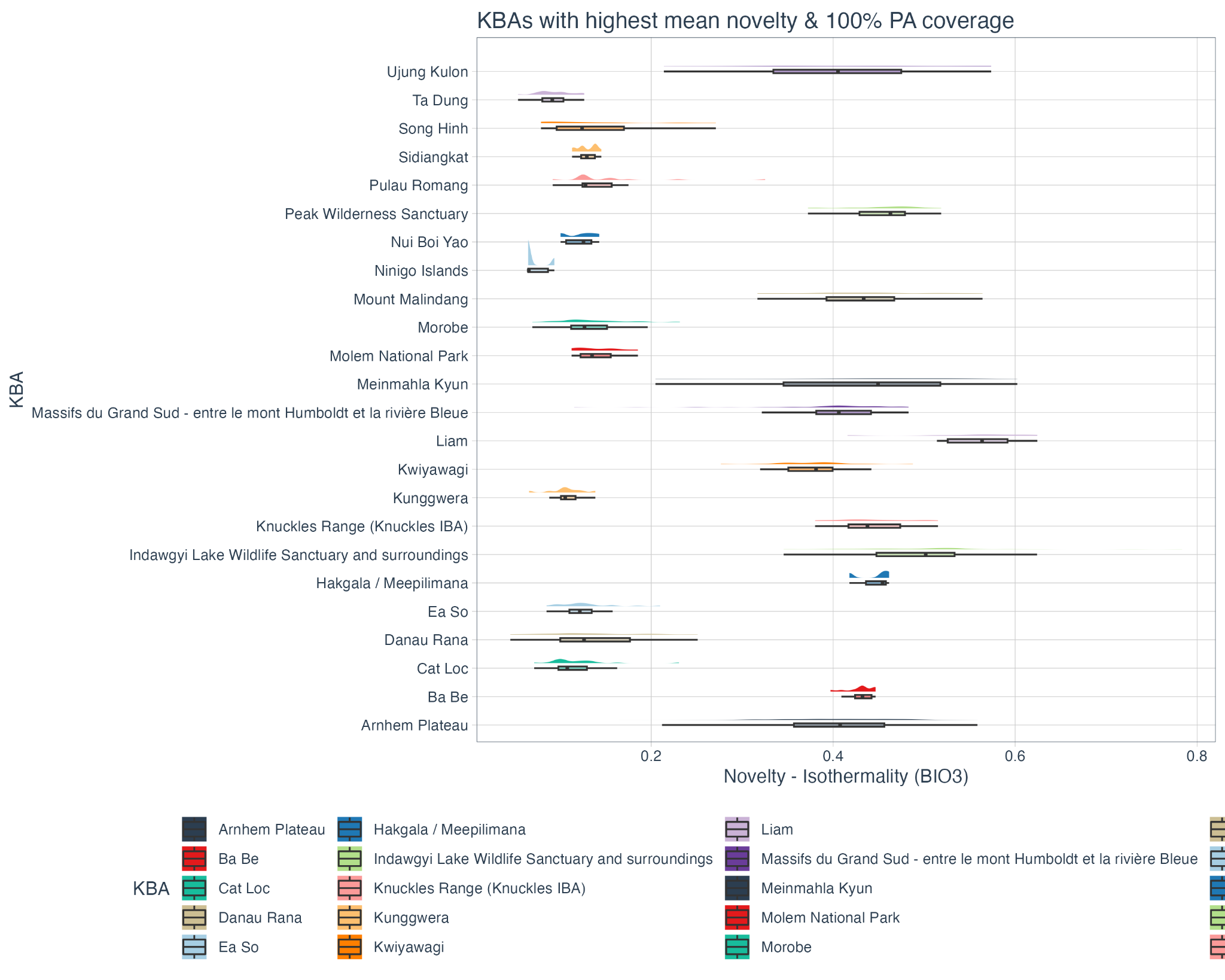

**Figure S4.6 |** Fractional novelty in recent isothermality (2005 to 2019) compared to the historic baseline (1990 to 2004) for the 12 KBAs in **Asia and Oceania** containing tropical forest with at least 99% protected area coverage and the highest recent mean fractional novelty in isothermality and the 12 KBAs in Asia and Oceania containing tropical forest with the lowest recent mean fractional novelty in isothermality and no protected area coverage. Novelty is measured between 0-1, where 1 indicates entirely novel isothermality regimes in 2005 - 2019.

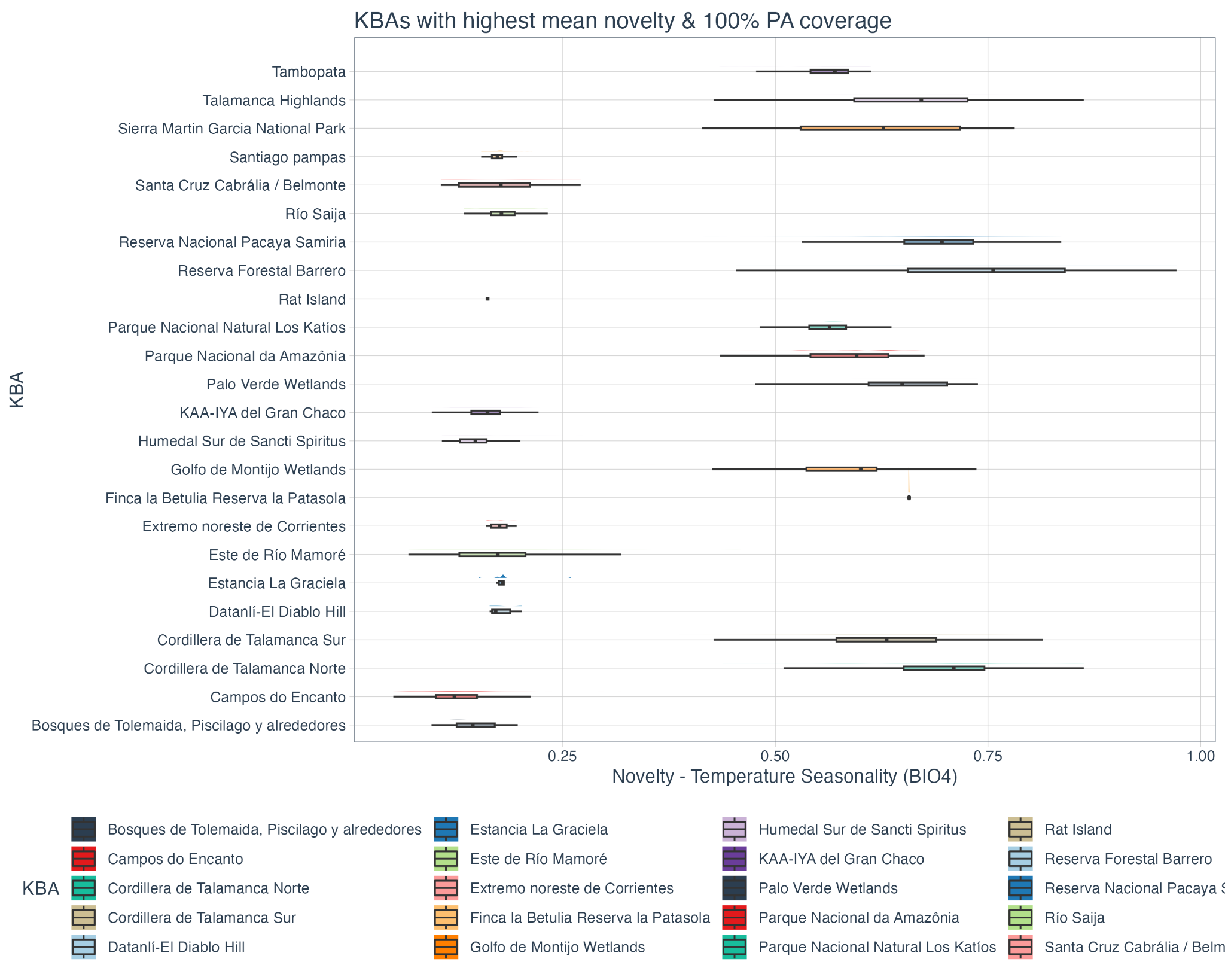

**Figure S4.7 |** Fractional novelty in recent temperature seasonality (2005 to 2019) compared to the historic baseline (1990 to 2004) for the 12 KBAs in **Latin America** containing tropical forest with at least 99% protected area coverage and the highest recent mean fractional novelty in temperature seasonality and the 12 KBAs in Latin America containing tropical forest with the lowest recent mean fractional novelty in temperature seasonality and no protected area coverage. Novelty is measured between 0-1, where 1 indicates entirely novel temperature seasonality regimes in 2005 - 2019.

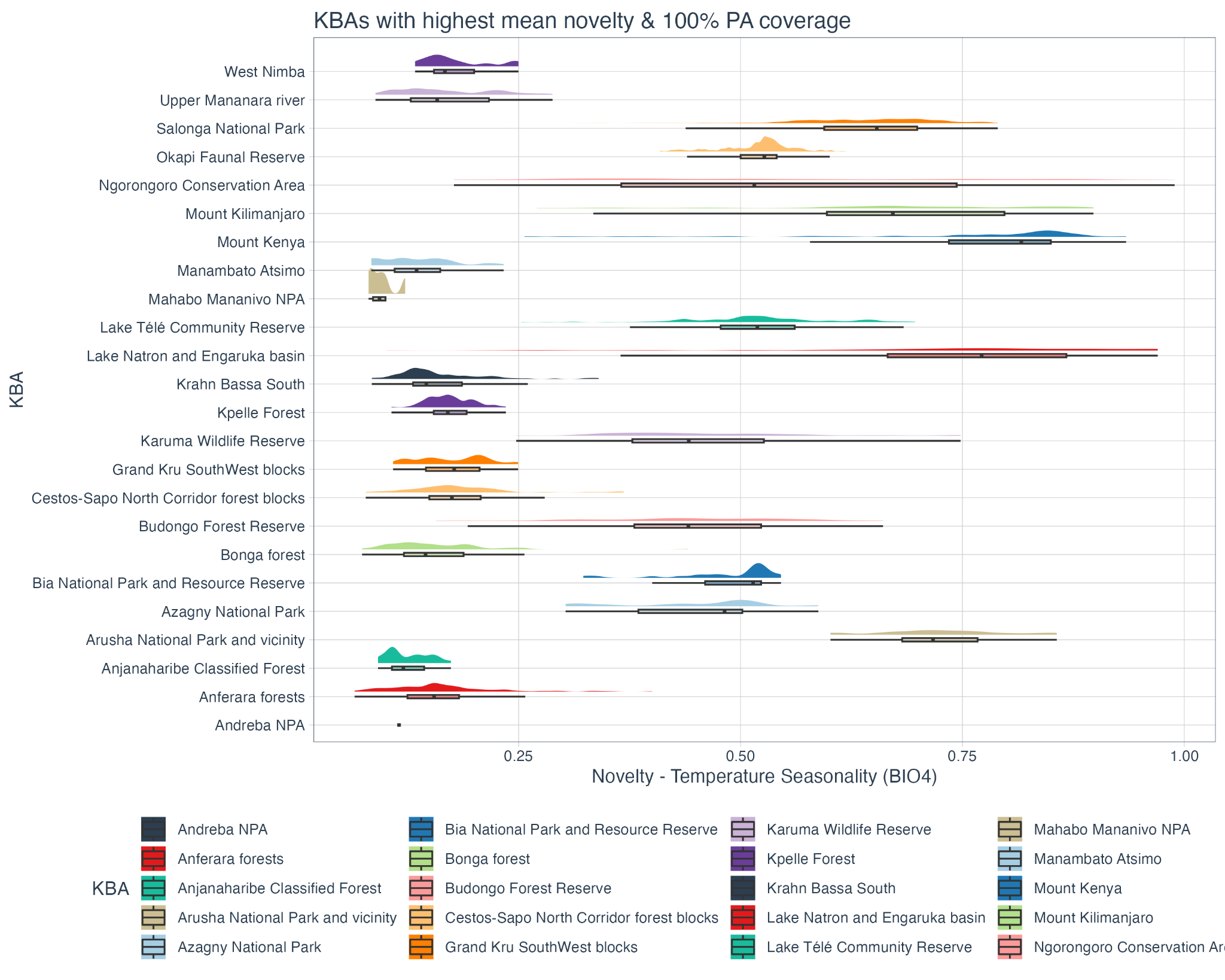

**Figure S4.8 |** Fractional novelty in recent temperature seasonality (2005 to 2019) compared to the historic baseline (1990 to 2004) for the 12 KBAs in **Africa** containing tropical forest with at least 99% protected area coverage and the highest recent mean fractional novelty in temperature seasonality and the 12 KBAs in Africa containing tropical forest with the lowest recent mean fractional novelty in temperature seasonality and no protected area coverage. Novelty is measured between 0-1, where 1 indicates entirely novel temperature seasonality regimes in 2005 - 2019.

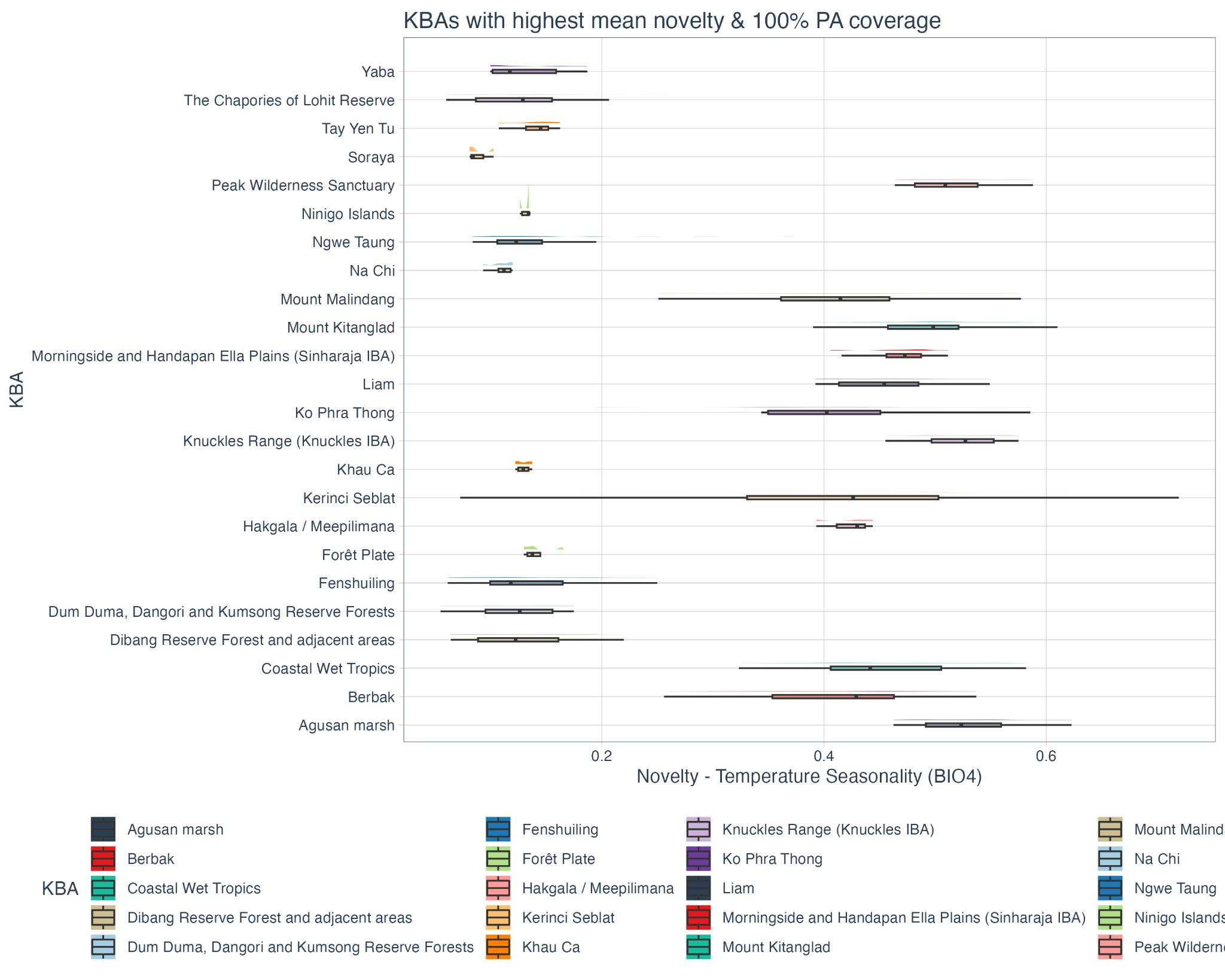

**Figure S4.9 |** Fractional novelty in recent temperature seasonality (2005 to 2019) compared to the historic baseline (1990 to 2004) for the 12 KBAs in **Asia and Oceania** containing tropical forest with at least 99% protected area coverage and the highest recent mean fractional novelty in temperature seasonality and the 12 KBAs in Asia and Oceania containing tropical forest with the lowest recent mean fractional novelty in temperature seasonality and no protected area coverage. Novelty is measured between 0-1, where 1 indicates entirely novel temperature seasonality regimes in 2005 - 2019.

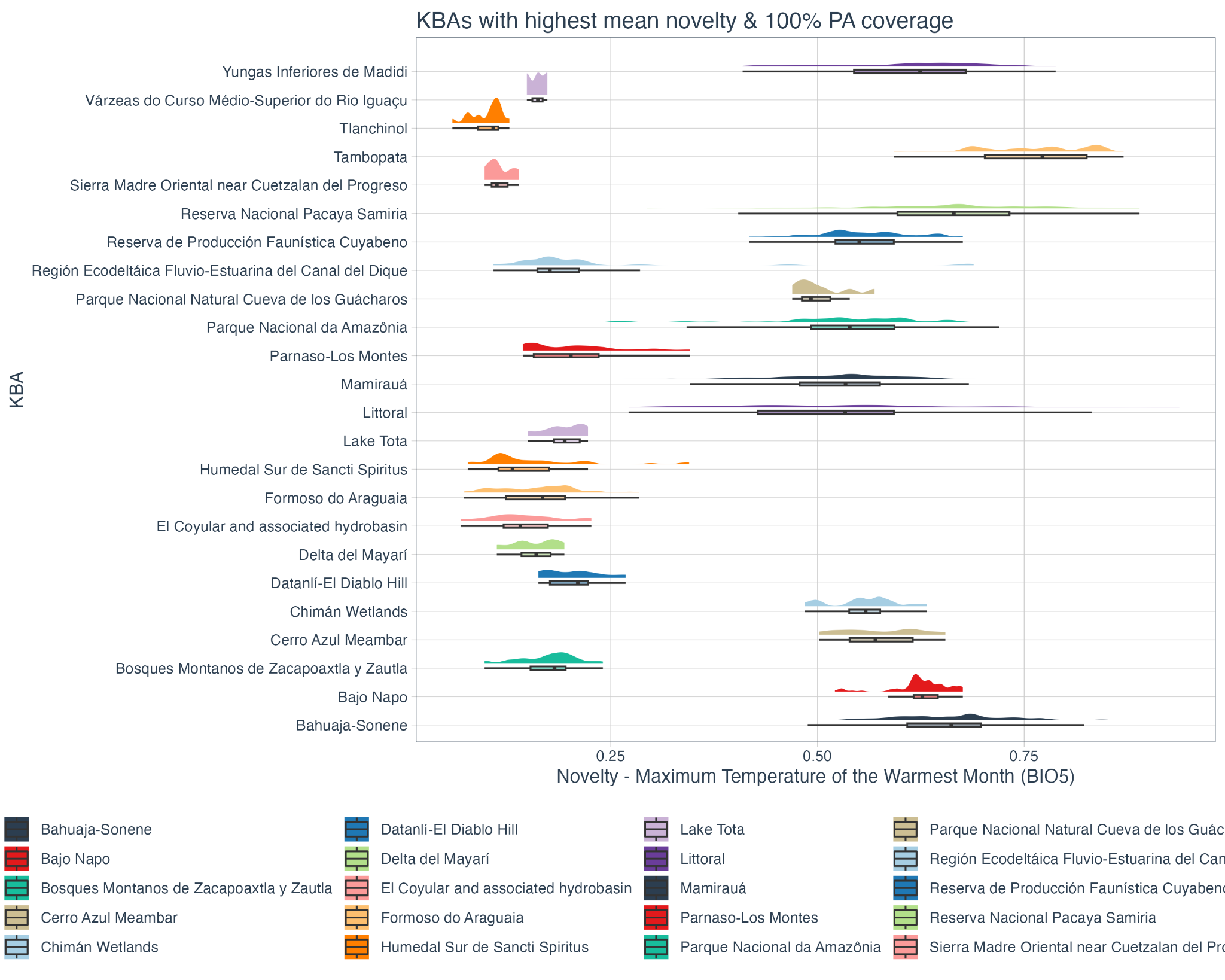

**Figure S4.10 |** Fractional novelty in recent maximum temperature of the warmest month (2005 to 2019) compared to the historic baseline (1990 to 2004) for the 12 KBAs in **Latin America** containing tropical forest with at least 99% protected area coverage and the highest recent mean fractional novelty in maximum temperature of the warmest month and the 12 KBAs in Latin America containing tropical forest with the lowest recent mean fractional novelty in maximum temperature of the warmest month and no protected area coverage. Novelty is measured between 0-1, where 1 indicates entirely novel maximum temperature of the warmest month regimes in 2005 - 2019.

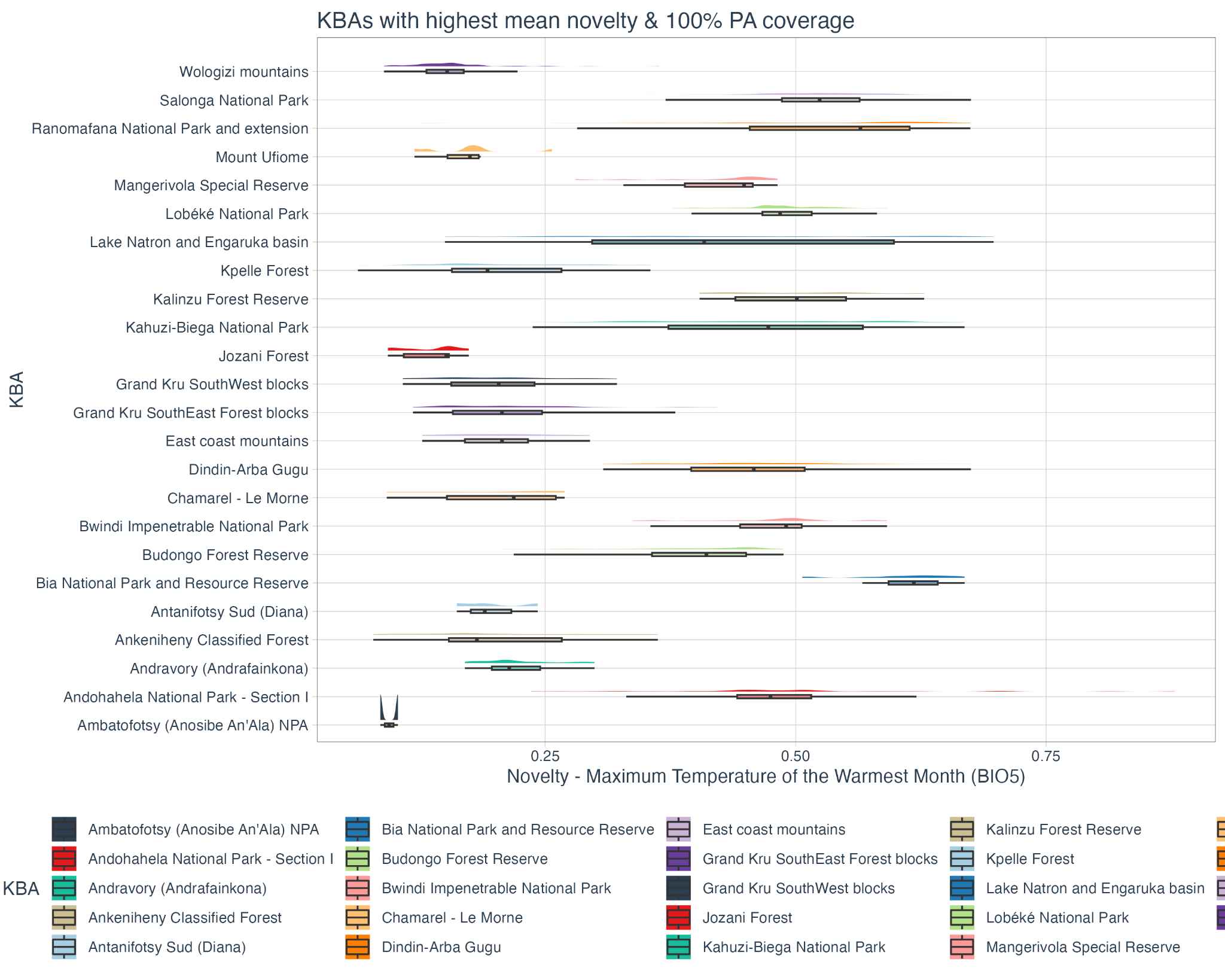

**Figure S4.11 |** Fractional novelty in recent maximum temperature of the warmest month (2005 to 2019) compared to the historic baseline (1990 to 2004) for the 12 KBAs in **Africa** containing tropical forest with at least 99% protected area coverage and the highest recent mean fractional novelty in maximum temperature of the warmest month and the 12 KBAs in Africa containing tropical forest with the lowest recent mean fractional novelty in maximum temperature of the warmest month and no protected area coverage. Novelty is measured between 0-1, where 1 indicates entirely novel maximum temperature of the warmest month regimes in 2005 - 2019.

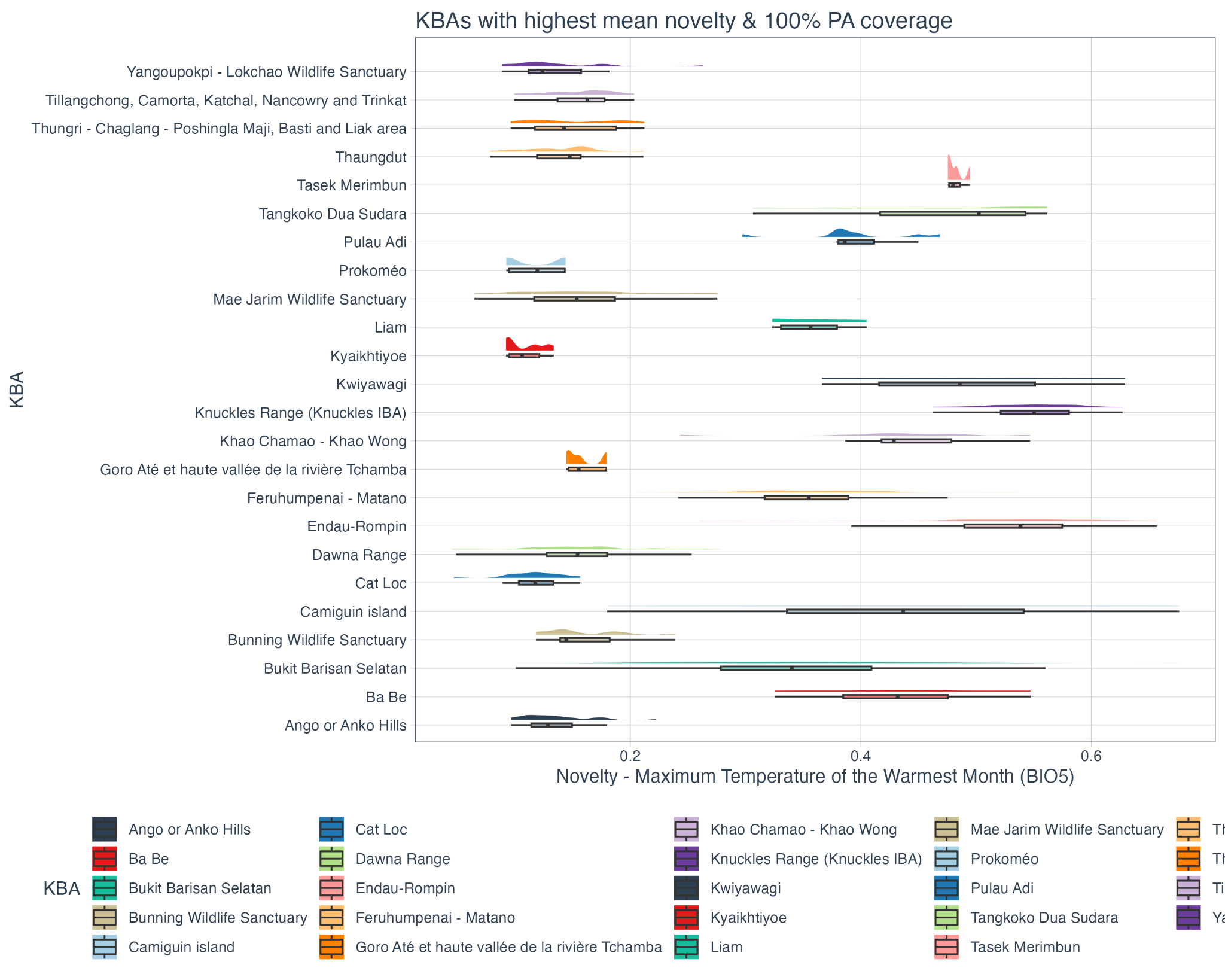

**Figure S4.12 |** Fractional novelty in recent maximum temperature of the warmest month (2005 to 2019) compared to the historic baseline (1990 to 2004) for the 12 KBAs in **Asia and Oceania** containing tropical forest with at least 99% protected area coverage and the highest recent mean fractional novelty in maximum temperature of the warmest month and the 12 KBAs in Asia and Oceania a containing tropical forest with the lowest recent mean fractional novelty in maximum temperature of the warmest month and no protected area coverage. Novelty is measured between 0-1, where 1 indicates entirely novel maximum temperature of the warmest month regimes in 2005 - 2019.

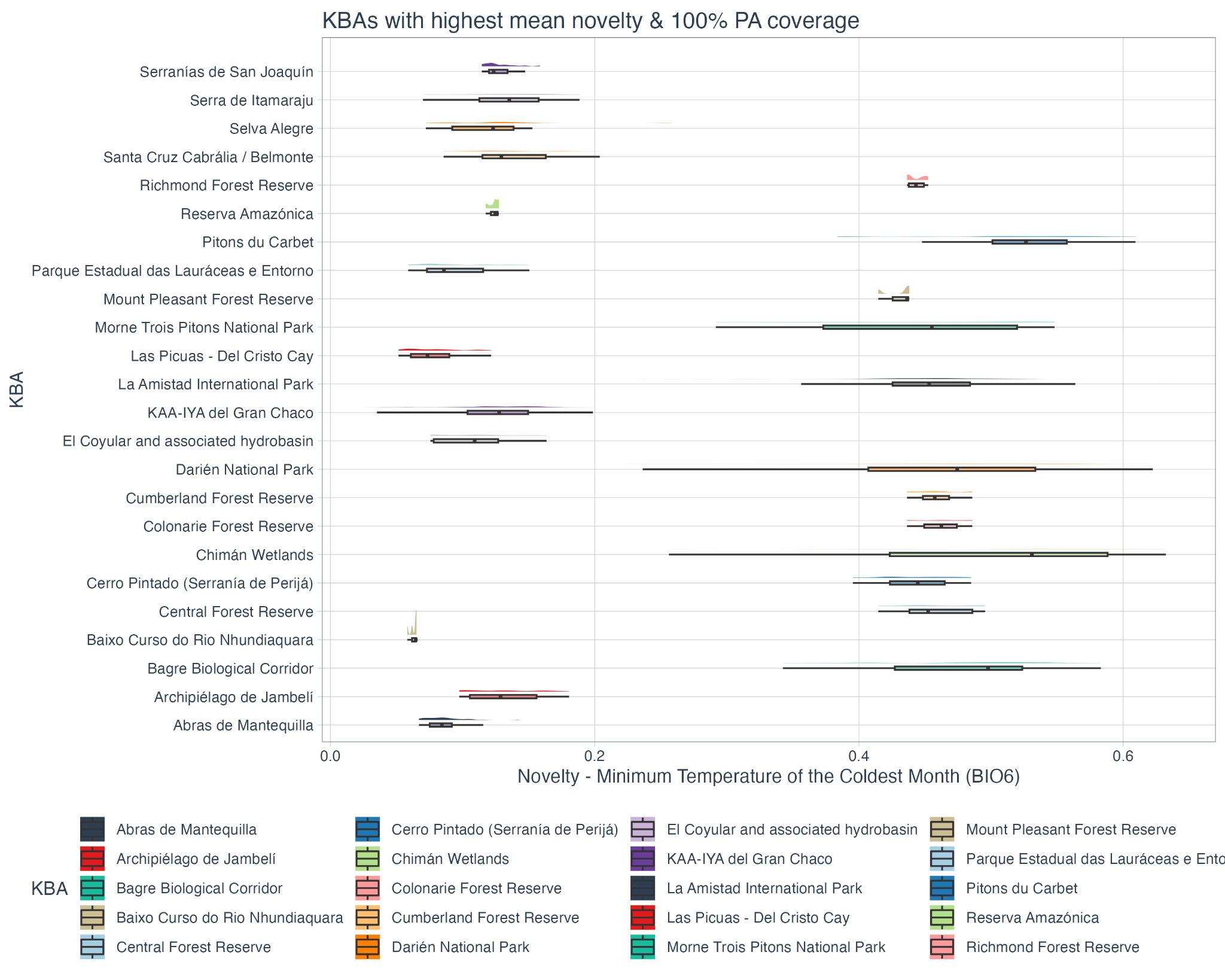

**Figure S4.13 |** Fractional novelty in recent minimum temperature of the coldest month (2005 to 2019) compared to the historic baseline (1990 to 2004) for the 12 KBAs in **Latin America** containing tropical forest with at least 99% protected area coverage and the highest recent mean fractional novelty in minimum temperature of the coldest month and the 12 KBAs in Latin America containing tropical forest with the lowest recent mean fractional novelty in minimum temperature of the coldest month and no protected area coverage. Novelty is measured between 0-1, where 1 indicates entirely novel minimum temperature of the coldest month regimes in 2005 - 2019.

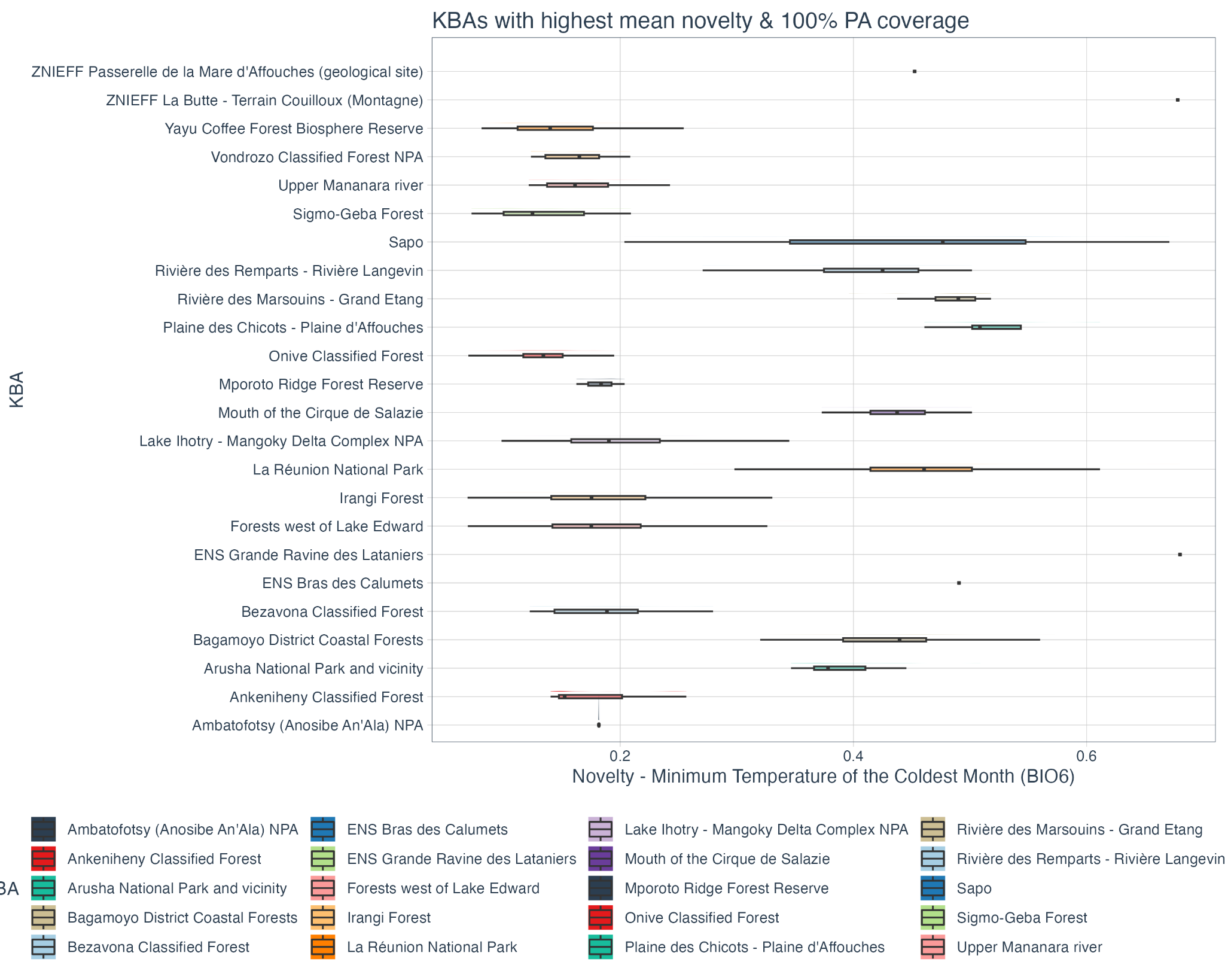

**Figure S4.14 |** Fractional novelty in recent minimum temperature of the coldest month (2005 to 2019) compared to the historic baseline (1990 to 2004) for the 12 KBAs in **Africa** containing tropical forest with at least 99% protected area coverage and the highest recent mean fractional novelty in minimum temperature of the coldest month and the 12 KBAs in Africa containing tropical forest with the lowest recent mean fractional novelty in minimum temperature of the coldest month and no protected area coverage. Novelty is measured between 0-1, where 1 indicates entirely novel minimum temperature of the coldest month regimes in 2005 - 2019.

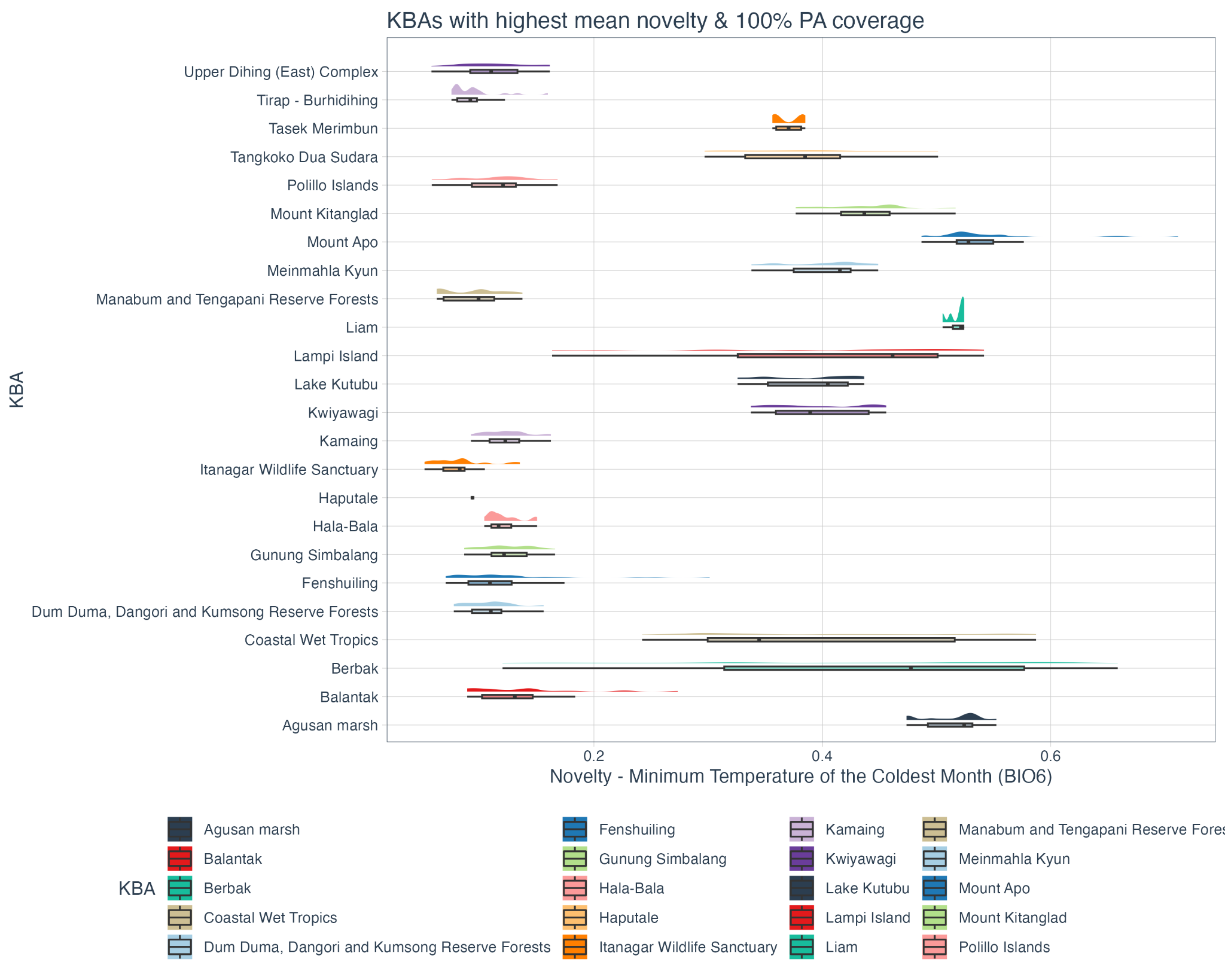

**Figure S4.15 |** Fractional novelty in recent minimum temperature of the coldest month (2005 to 2019) compared to the historic baseline (1990 to 2004) for the 12 KBAs in **Asia and Oceania** containing tropical forest with at least 99% protected area coverage and the highest recent mean fractional novelty in minimum temperature of the coldest month and the 12 KBAs in Asia and Oceania containing tropical forest with the lowest recent mean fractional novelty in minimum temperature of the coldest month and no protected area coverage. Novelty is measured between 0-1, where 1 indicates entirely novel minimum temperature of the coldest month regimes in 2005 - 2019.

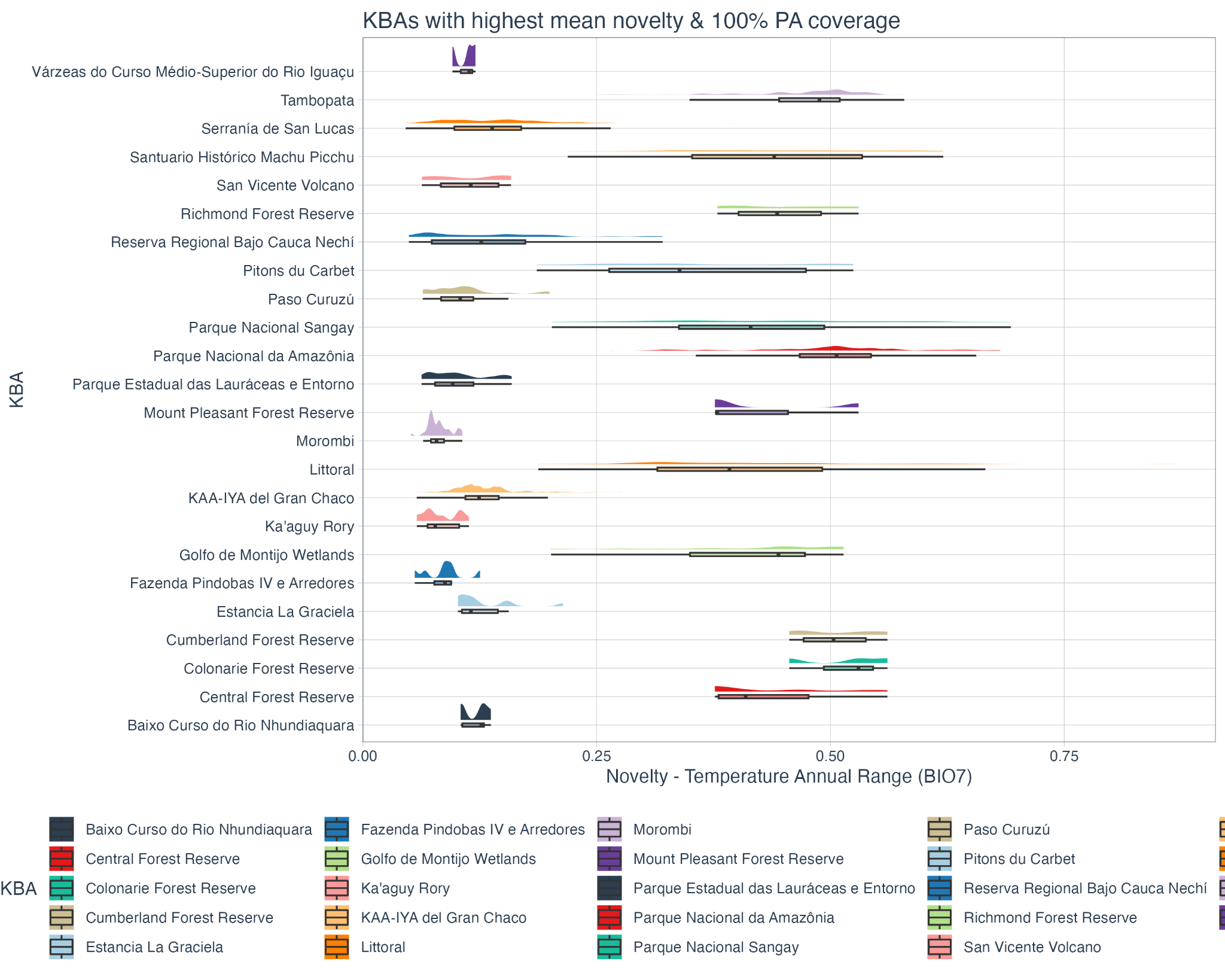

**Figure S4.16 |** Fractional novelty in recent annual temperature range (2005 to 2019) compared to the historic baseline (1990 to 2004) for the 12 KBAs in **Latin America** containing tropical forest with at least 99% protected area coverage and the highest recent mean fractional novelty in annual temperature range and the 12 KBAs in Latin America containing tropical forest with the lowest recent mean fractional novelty in annual temperature range and no protected area coverage. Novelty is measured between 0-1, where 1 indicates entirely novel annual temperature range regimes in 2005 - 2019.

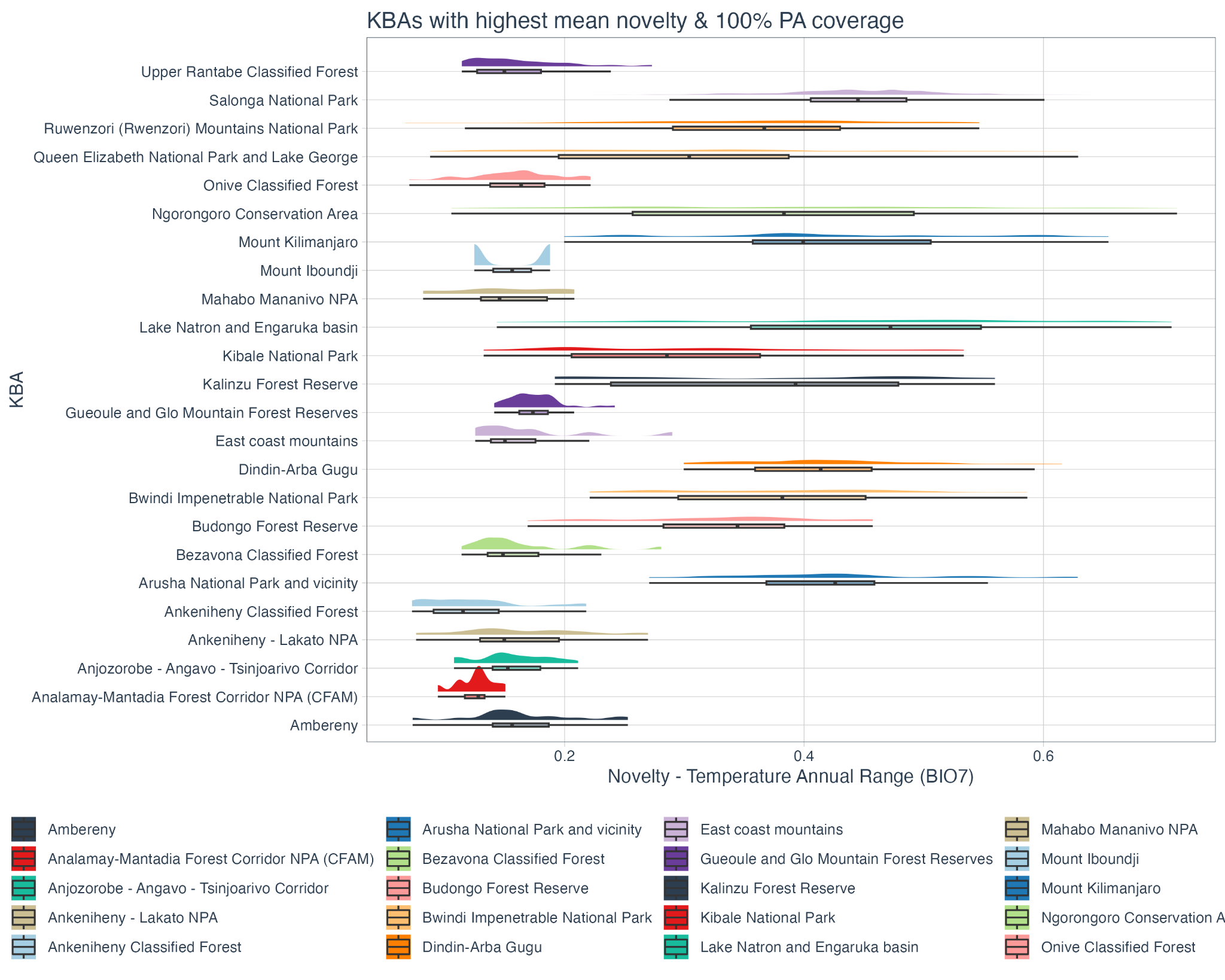

**Figure S4.17 |** Fractional novelty in recent annual temperature range (2005 to 2019) compared to the historic baseline (1990 to 2004) for the 12 KBAs in **Africa** containing tropical forest with at least 99% protected area coverage and the highest recent mean fractional novelty in annual temperature range and the 12 KBAs in Africa containing tropical forest with the lowest recent mean fractional novelty in annual temperature range and no protected area coverage. Novelty is measured between 0-1, where 1 indicates entirely novel annual temperature range regimes in 2005 - 2019.

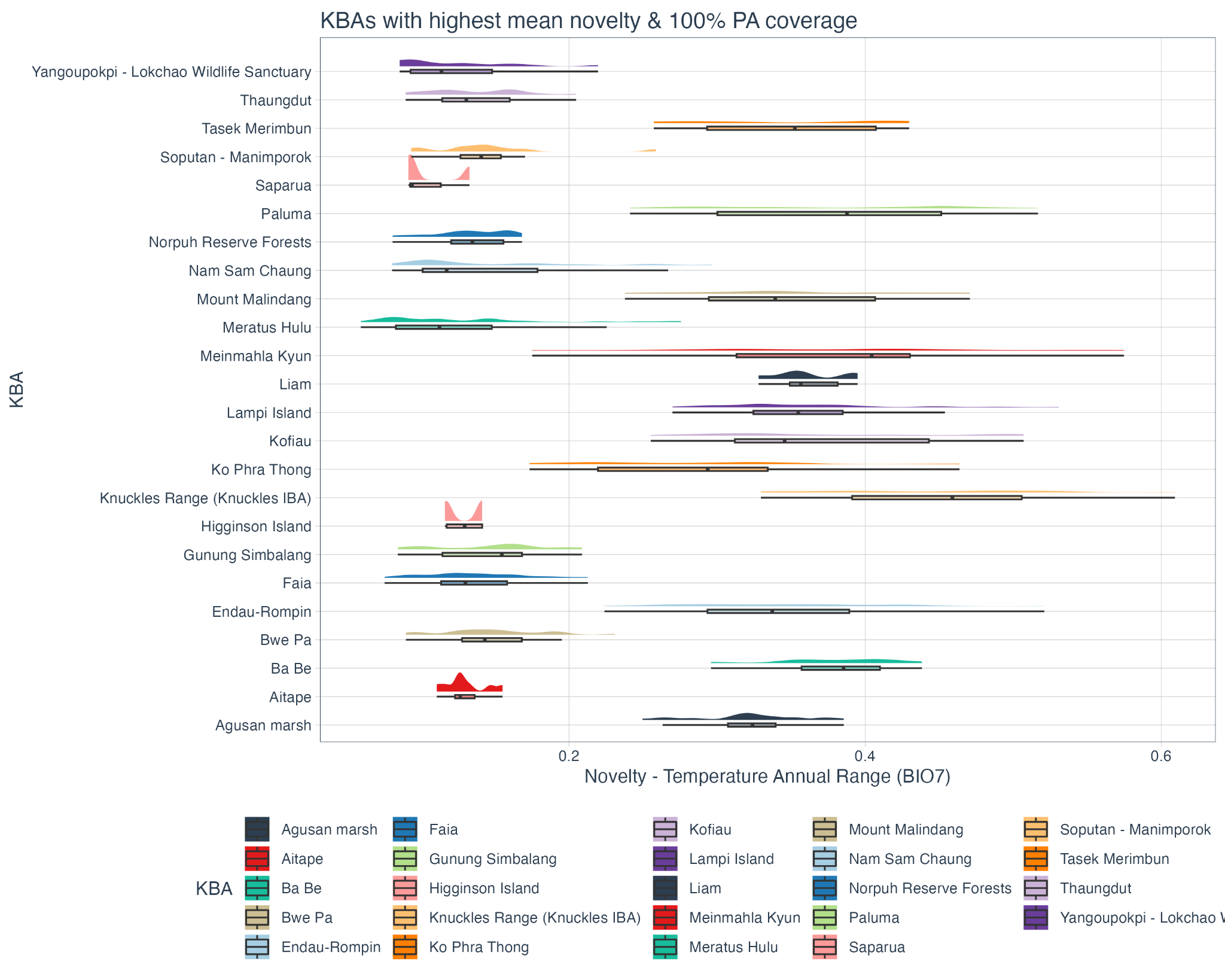

**Figure S4.18 |** Fractional novelty in recent annual temperature range (2005 to 2019) compared to the historic baseline (1990 to 2004) for the 12 KBAs in **Asia and Oceania** containing tropical forest with at least 99% protected area coverage and the highest recent mean fractional novelty in annual temperature range and the 12 KBAs in Asia and Oceania containing tropical forest with the lowest recent mean fractional novelty in annual temperature range and no protected area coverage. Novelty is measured between 0-1, where 1 indicates entirely novel annual temperature range regimes in 2005 - 2019.
